# scVAE: Variational auto-encoders for single-cell gene expression data

**DOI:** 10.1101/318295

**Authors:** Christopher Heje Grønbech, Maximillian Fornitz Vording, Pascal Timshel, Casper Kaae Sønderby, Tune Hannes Pers, Ole Winther

**Author notes:** The length of this alphabet is 127.55219pt.

## Abstract

**Motivation:** Models for analysing and making relevant biological inferences from massive amounts of complex single-cell transcriptomic data typically require several individual data-processing steps, each with their own set of hyperparameter choices. With deep generative models one can work directly with count data, make likelihood-based model comparison, learn a latent representation of the cells and capture more of the variability in different cell populations.

**Results:** We propose a novel method based on variational auto-encoders (VAEs) for analysis of single-cell RNA sequencing (scRNA-seq) data. It avoids data preprocessing by using raw count data as input and can robustly estimate the expected gene expression levels and a latent representation for each cell. We tested several count likelihood functions and a variant of the VAE that has a priori clustering in the latent space. We show for several scRNA-seq data sets that our method outperforms recently proposed scRNA-seq methods in clustering cells and that the resulting clusters reflect cell types.

**Availability and implementation:** Our method, called scVAE, is implemented in Python using the TensorFlow machine-learning library, and it is freely available at https://github.com/scvae/scvae.

## 1 Introduction

Single-cell RNA sequencing (scRNA-seq) enables measurement of gene expression levels of individual cells and thus provides a new framework to understand dysregulation of disease at the cell-type level (Regev *et al.*, 2017). To date, a number of methods^1^ have been developed to process gene expression data to normalise the data (Haghverdi *et al.*, 2018) and cluster cells into putative cell types (Satija *et al.*, 2015; Kiselev *et al.*, 2017). Seurat (Satija *et al.*, 2015) is a popular method which is a multi-step process of normalisation, transformation, decomposition, embedding, and clustering of the gene expression levels. This can be cumbersome, and a more automated approach is desirable. Four recent methods, cellTree (duVerle *et al.*, 2016), DIMM-SC (Sun *et al.*, 2017), DCA (Eraslan *et al.*, 2018), and scVI (Lopez *et al.*, 2018), model the gene expression levels directly as counts using latent Dirichlet allocation, Dirichlet-mixture generative model, an auto-encoder, and a variational auto-encoder, respectively. The last two methods are described below.

Here, we show that expressive deep generative models, leveraging the recent progress in deep neural networks, provide a powerful framework for modelling the data distributions of raw count data. We show that these models can learn biologically plausible representations of scRNA-seq experiments using only the highly sparse raw count data as input entirely skipping the normalisation and transformation steps needed in previous methods. Our approach is based on the variational auto-encoder (VAE) framework presented in Kingma and Welling (2013) and Rezende *et al.* (2014). These models learn compressed latent representations of the input data without any supervisory signals, and they can denoise the input data using its encoder-decoder structure. We extend these models with likelihood (link) functions suitable for modelling (sparse) count data and perform extensive experiments on several data sets to analyse the strengths and weaknesses of the models and the likelihood functions. Variational auto-encoders have been examined and extended to use multiple latent variables for, e.g., semi-supervised learning (Kingma *et al.*, 2014), clustering (Dilokthanakul *et al.*, 2016; Maaløe *et al.*, 2017; Jiang *et al.*, 2017), and structured representations (Sønderby *et al.*, 2016; Johnson *et al.*, 2016; Lin *et al.*, 2018). They have also been used in variety of cases to, e.g., generate sentences (Bowman *et al.*, 2016), transfer artistic style of paintings (Gatys *et al.*, 2015), and create music (Roberts *et al.*, 2017). Recently, VAEs have also been used to study normalised bulk RNA-seq data (Way and Greene, 2017) as well as visualising normalised scRNA-seq data (VASC, Wang and Gu, 2018; scvis, Ding *et al.*, 2018).

Compared to most other auto-encoder methods, VAEs have two major advantages: (a) It is probabilistic so that performance can be quantified and compared in terms of the likelihood, and (b) the variational objective creates a natural trade-off between data reconstruction and model capacity. Other types of auto-encoders also learn a latent representation from data that is used to reconstruct the same as well as unseen data. The latent representation is tuned to have low-capacity, so the model is forced to only estimate the most important features of the data. Different auto-encoder models have previously been used to model normalised (or binarised) bulk gene expression levels: denoising auto-encoders (Tan *et al.*, 2014; Gupta *et al.*, 2015; Tan *et al.*, 2016), sparse auto-encoders (Chen *et al.*, 2016), and robust auto-encoders (Cui *et al.*, 2017). A bottleneck auto-encoder (Eraslan *et al.*, 2018) was also recently used to model single-cell transcript counts, and a generative adversarial network (Goodfellow *et al.*, 2014), which is a related model, was recently used to model normalised single-cell gene expression levels (Ghahramani *et al.*, 2018).

Our contributions are threefold: (a) We have developed new generative models based on the VAE framework for directly modelling raw counts from RNA-seq data; (b) we show that our models using either a Gaussian or a Gaussian-mixture latent variable prior learn biologically plausible groupings of scRNA-seq data of as validated on data sets with known cell types; and (c) we provide a publicly available framework for unsupervised modelling of count data from RNA-seq experiments.

## 2 Methods and materials

We have developed generative models for directly modelling the raw read counts from scRNA-seq data. In this section, we describe the models as well as the different data likelihood (link) functions for this task.

### 2.1 Latent-variable models

We take a generative approach to modelling raw count data vectors **x**, where **x** represents a single cell and its components *x*_*n*_, which are also called *features*, correspond to the gene expression count for gene *n*. We assume that **x** is drawn from the distribution *p*_***θ***_(**x**) = *p*(**x** | ***θ***) parameterised by ***θ***. The number of features (genes), *N*, is very high, but we still want to model co-variation between the features. We achieve this by introducing a stochastic latent variable **z**, with fewer dimensions than **x**, and condition the data-generating process on this. The joint probability distribution of **x** and **z** is then

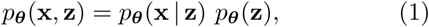

where *p*_***θ***_(**x** | **z**) is the likelihood function and *p*_***θ***_(**z**) is the prior probability distribution of **z**. Marginalising over **z** results in the marginal likelihood of ***θ*** for one data point:

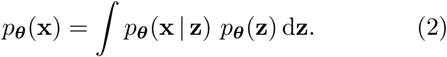

The log-likelihood function for all data points (also called *examples*, which in this case are cells) will be the sum of the log-likelihoods for each data point: 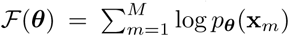, where *M* is the number of cells. We can then use maximum-likelihood estimation, arg max_***θ***_ ***ℱ*** (***θ***), to infer the value ***θ***.

We use a deep neural network to map from **z** to the sufficient statistics of **x**. However, as the marginalisation over the latent variables in Equation (2) is intractable for all but the simplest linear models, we have to resort to approximate methods for inference in the models. Here we use variational Bayesian optimisation for inference as introduced in the variational auto-encoder framework (Kingma and Welling, 2013; Rezende *et al.*, 2014). Details of this approach is covered in Section 2.4.

### 2.2 Deep generative models

The choice of likelihood function *p*_***θ***_(**x** | **z**) depends on the statistical properties of the data, e.g., continuous or discrete observations, sparsity, and computational tractability. Contrary to most other VAE-based articles considering either continuous or categorical input data, our goal is to model discrete count data directly, which is why Poisson or negative binomial likelihood functions are natural choices. This will be discussed in more detail below. The prior over the latent variables *p*(**z**) is usually chosen to be an isotropic standard multivariate Gaussian distribution to get the following generative process (Fig. 1A):

**Fig. 1.**
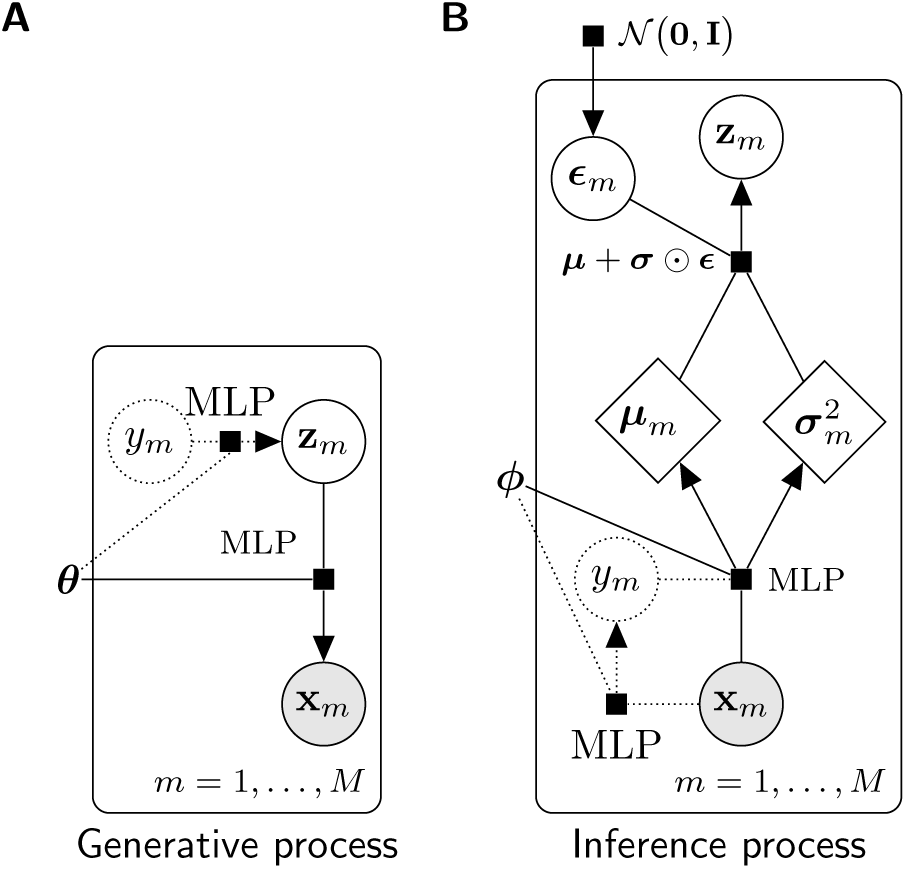
Model graphs of (**A**) the generative process and (**B**) the inference process of the variational auto-encoder models. Dotted strokes designate nodes and edges for the Gaussian-mixture model only. Grey circles signify observable variables, white circles represent latent variables, and rhombi symbolise deterministic variables. The black squares denote the functions next to them with the variables connected by lines as input and the variable connected by an arrow as output.

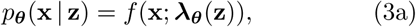

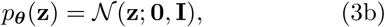

where *f* is a discrete distribution such as the Poisson (P) distribution: 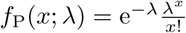 and *f* (**x**; ***λ***) = Π_*k*_ *f* (*x*_*k*_; *λ*_*k*_). The Poisson rate parameters are functions of **z**: ***λ***_***θ***_(**z**). We can for example parameterise it by a single-layer feed-forward neural network:

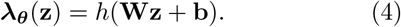

Here, **W** and **b** are the weights and bias of a linear model, ***θ*** = (**W, b**), and *h*(·) is an appropriate element-wise non-linear transformation to make the rate non-negative. The Poisson likelihood function can be substituted by other probability distributions with parameters of the distribution being nonlinear functions of **z** in the same fashion as above (see Section 2.3). In order to make the model more expressive, we can also replace the single-layer model with a deep model with one or more hidden layers. For *L* layers of adaptable weights we can write: **a**_*l*_ = *h*_*l*_(**W**_*l*_**a**_*l*−1_ + **b**_*l*_) for *l* = 1, …, *L*, **a**_0_ = **z**, and ***λ***_***θ***_(**z**) = **a**_*L*_ with *h*_*l*_(·) denoting the activation function of the *l*th layer. For the hidden layers the rectifier function, ReLU(*x*) = max(0, *x*), is often a good choice.

#### 2.2.1 Gaussian-mixture variational auto-encoder

Using a Gaussian distribution as the prior probability distribution of **z** only allows for one mode in the latent representation. If there is an inherent clustering in the data, like for scRNA-seq data where the cells represent different cell types, it is desirable to have multiple modes, e.g., one for every cluster or class. This can be implemented by using a Gaussian-mixture model in place of the Gaussian distribution. Following the specification of the M2 model of Kingma *et al.* (2014), with inspiration from Dilokthanakul *et al.* (2016) and modifications by Rui Shu^2^, a categorical latent random variable *y ∈*{1, …, *K*} is added to the VAE model:

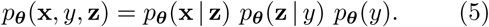

where the likelihood term *p*_***θ***_(**x** | **z**) is same as for the standard VAE (Eq. 3a) and

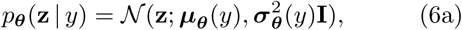

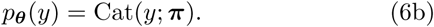

The generative process for the Gaussian-mixture variational auto-encoder (GMVAE) is illustrated in Figure 1A. Here, ***π*** is a *K*-dimensional probability vector, where *K* is the number of components in the Gaussian-mixture model. The component *π*_*k*_ of ***π*** is the mixing coefficient of the *k*th Gaussian distribution, quantifying how much this distribution contributes to the overall probability distribution. Without any prior knowledge, the categorical distribution is fixed to be uniform.

### 2.3 Modelling gene expression count data

Instead of normalising and transforming the gene expression data, the original transcript counts are modelled directly to take into account the total amount of genes expressed in each cell also called the *sequencing depth*. To model count data the likelihood function *p*_***θ***_(**x** | **z**) will need to be discrete and only have non-negative support. As described earlier, parameters for these probability distributions are modelled using deep neural networks that takes the latent vector **z** as input. We will consider a number of such distributions in the following and investigate which ones are best in term of likelihood on held-out data.

Our first approach to model the likelihood function uses either the Poisson or negative binomial (NB) distributions. The Poisson distribution provides the simplest distribution naturally handling count data, whereas the negative binomial distribution corresponds well with the over-dispersed nature of gene expression data (Oshlack *et al.*, 2010). To properly handle the sparsity of the scRNA-seq data (Vallejos *et al.*, 2017), we test two approaches: a zero-inflated distribution and modelling of low counts using a discrete distribution. A zero-inflated distribution adds an additional parameter, which controls the amount of excessive zeros added to an existing probability distribution. For the Poisson distribution, *f*_P_(*x*; *λ*), the zero-inflated version (ZIP) is defined as

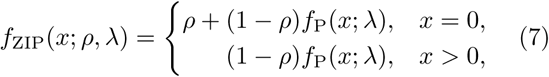

where *ρ* is the probability of excessive zeros. The zero-inflated negative binomial (ZINB) distribution have an analogous expression.

The so-called piece-wise categorical distribution, which has been used for demand forecasting (Seeger *et al.*, 2016), uses a mixture of discrete probabilities for low counts and a shifted probability distribution for counts equal to or greater than *k*_max_. The piece-wise categorical version of the Poisson distribution (PCP) is

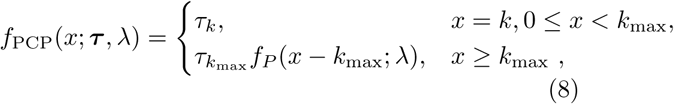

where *τ*_*k*_, *k* = 0, …, *k*_max_ − 1, is the probability of a count being equal to *k* and 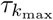 the probability that the count is *k*_max_ or above.

Lastly, we also investigate the so-called constrained Poisson (CP) distribution (Salakhutdinov and Hinton, 2009) which reparameterises the Poisson to depend explicitly on the sequencing depth *D*_*m*_ of cell *m*. For the constrained Poisson distribution, the rate parameter *λ*_*mn*_ for gene *n* in cell *m* is

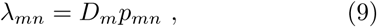

where *p*_*mn*_ is constrained to be a probability distribution for each cell: *p*_*mn*_ ≥ 0 and Σ_*n*_ *p*_*mn*_ = 1. This construction ensures the rates are set such that the expected number of counts are equal to the sequencing depth while we can still model each gene by a Poisson distribution.

Since we do not know the true conditioning structure of the genes, we make the simplifying assumption that they are independent for computational reasons and therefore use feed-forward neural networks.

### 2.4 Variational auto-encoders

In order to train our deep generative models, we will use the variational auto-encoder framework which amounts to replacing the intractable marginal likelihood with its variational lower bound and estimate the intractable integrals with low-variance Monte Carlo estimates. The lower bound is maximised for the training set and then evaluated on a test set in order to perform model comparison.

Since the likelihood function *p*_***θ***_(**x** | **z**) is modelled using non-linear transformations, the posterior probability distribution *p*_***θ***_(**z** | **x**) = *p*_***θ***_(**x** | **z**)*p*_***θ***_(**z**)*/p*_***θ***_(**x**) becomes intractable. Variational auto-encoders use a variational Bayesian approach where *p*_***θ***_(**z** | **x**) is replaced by an approximate probability distribution *q*_***ϕ***_(**z** | **x**) modelled using non-linear transformations parameterised by ***ϕ***. Thus, the marginal log-likelihood can be written as

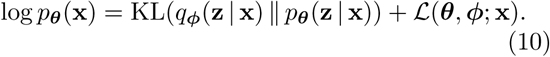

Here, the first term is the Kullback–Leibler (KL) divergence between the true and the approximate posterior distributions. ℒ (***θ, ϕ***) can be rewritten as

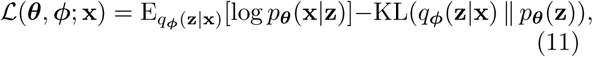

where the first term is the *expected negative reconstruction error* quantifying how well the model can reconstruct **x**, and the second term is the *relative KL divergence* between the approximate posterior distribution *q*_***ϕ***_(**z**|**x**) and the prior distribution *p*_***θ***_(**z**) of **z**.

Evaluating Equation (10) is intractable due to the appearance of the intractable true posterior distribution. However, as the KL divergence is non-negative, Equation (11) provides a lower bound on the marginal, e.g., log *p*_***θ***_(**x**) ≥ ℒ (***θ, ϕ***; **x**). The integrals over **z** in Equation (11) are still analytically intractable, but a low-variance estimator can be constructed using Monte Carlo integration as mentioned above.

For the standard VAE, the approximate posterior distribution is chosen to be a multivariate Gaussian distribution with a diagonal covariance matrix:

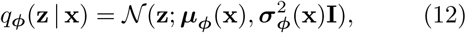

where ***µ***_***ϕ***_(**x**) and 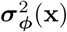 are non-linear transformations of **x** parameterised by a neural network with parameters ***ϕ*** in a similar fashion as Equation (4) is used to parameterise the generative model. Equation (12) is called the inference model, and it is illustrated in Figure 1B together with the inference model for the GMVAE, which is described in Supplementary Section S1.

The parameters of the generative and inference models are optimised simultaneously using a stochastic gradient descent algorithm. The reparameterisation trick for sampling from *q*_***ϕ***_(**z** | **x**) allows back-propagation of the gradients end-to-end (Kingma and Welling, 2013). We use a *warm-up* scheme during optimisation as described in the Supplementary Section S2, and we report the marginal log-likelihood lower bound averaged over all examples. The VAE *reconstruction* of **x** is defined as the mean of the expression values using a stochastic encoding and decoding step: 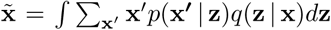. The variance of the reconstruction can also be computed in a similar fashion.

### 2.5 Clustering and visualisation of the latent space

The aim of using a VAE is to capture the joint distribution of multivariate data such as gene expression profiles. Since our method is likelihood-based, model comparison can be performed by evaluating the marginal log-likelihood (lower bound) on a test set for different models. For model comparison with nonlikelihood-based methods or methods that preprocess the data, we need other metrics such as a clustering quality. For latent variable methods, clustering in the latent space is the most obvious approach. Therefore, we do this as well.

The categorical latent variable used in the GMVAE model gives a built-in clustering by using as assignment the mixing coefficient with the highest *responsibility q*_***ϕ***_(*y* | **x**). The responsibilities are computed as part of the inference network as described in Supplementary Section S1. The standard VAE model does not have this feature built in, so instead *k*-means clustering (kM) is used to cluster the latent representations of the cells for this model. We also test *k*-means clustering for the GMVAE model.

We use two clustering metrics to measure the similarity between a clustering found by one of our models and the clustering given in the data set. These are the adjusted Rand index (Hubert and Arabie, 1985), *R*_adj_, and adjusted mutual information (AMI; Vinh *et al.* (2009)). For both metrics, two identical clusterings have a value of 1, while the expected value of the metrics is 0 (they are not bounded below).

To visualise the latent space in which the latent representations reside, the mean values of the approximate posterior distribution *q*_***ϕ***_(**z** | **x**) are plotted instead of the samples. This is done to better get a representation of the probability distribution for the latent representation of each cell. For the standard VAE, this is just 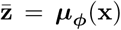, and for the GMVAE, 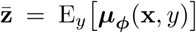. For latent spaces with only two dimensions, we plot these directly, but to visualise higher dimensional latent spaces, we project to two dimensions using *t*-distributed stochastic neighbour embeddings (*t*-SNE; van der Maaten and Hinton, 2008).

### 2.6 RNA-seq data sets

We model four different RNA-seq data sets as summarised in Table 1. The first data set is of single-cell gene expression counts of peripheral blood mononuclear cells (PBMC) used for generation of reference transcriptome profiles (Zheng *et al.*, 2017). It is combined from nine separate data sets of different purified cell types and published by 10x Genomics^3^, and we use the filtered gene–cell matrices for each data set. Another data set of purified monocytes is also available, but since another cell type was discovered in this data set (Zheng *et al.*, 2017), and since no separation of the two is available in the data set, we only use the other nine data sets, which are listed in Table 1. This table also shows the high degree of sparsity of the data set. The second data set is a bulk RNA-seq data set made publicly available by TCGA (Weinstein *et al.*, 2013).^4^ This data set consists of RSEM (Li and Dewey, 2011) expected gene expression counts for samples of human cancer cells from 29 tissue sites. Gene IDs are used in this data set, so the available mapping from TCGA is used to map the IDs to gene names. The difference in number of genes and sparsity from the original data set is not large. The two remaining data sets are described, modelled, and analysed in Supplementary Sections S4–5.

**Table 1.** Overview of gene expression data sets.

| Data set | Examples | Features | Classes | Sparsity |
| --- | --- | --- | --- | --- |
| PBMC | 92 043 | 32 738 | 9<sup>a</sup> | 0.980 |
| MBC | 1 306 127 | 27 998 | — | 0.928 |
| TCGA | 10 830 | 58 581 | 29 | 0.525 |
| GTE<sub>x</sub> | 11 688 | 54 271 | 53 | 0.472 |

<sup>a</sup> CD19+ B cells, CD34+ cells, CD4+ helper T cells, CD4+/CD25+ regulatory T cells, CD4+/CD45RA+/CD25-naïve T cells, CD4+/CD45RO+ memory T cells, CD56+ natural killer cells, CD8+ cytotoxic T cells, CD8+/CD45RA+ naïve cytotoxic T cells.

### 2.7 Experiment setup

Each data set is divided into training, validation, and test sets using a 81 %–9 %–10 % split with uniformly random selection. The training sets are used to train the models, the validation sets are used for hyperparameter selection during training, and the test sets are used to evaluate the models after training.

For the deep neural networks, we examine different network architectures to find the optimal one for each data set. We test deep neural networks of both one and two hidden layers with 100, 250, 500, or 1000 units each. We also experiment with a latent space of both 10, 25, 50, and 100 dimensions. A standard VAE with the negative binomial distribution as the likelihood function (a VAE-NB model) is used for these experiments. Using the optimal architecture, we test the link functions introduced in Section 2.3 as likelihood function for both the standard VAE as well as the GMVAE. We train and evaluate each model three times for each likelihood function.

The hidden units in the deep neural networks use the rectifier function as their non-linear transformation. For real, positive parameters (*λ* for the Poisson distribution, *r* for the negative binomial distribution, *σ* in the Gaussian distribution), we model the natural logarithm of the parameter. For the standard deviation *σ* in the Gaussian-mixture model, however, we use the softplus function, log(1 + e^*x*^), to constrain the possible covariance matrices to be only positive-definite. The units modelling the probability parameters in the negative binomial distribution, *p*, and the zero-inflated distributions, *π*_*k*_, use the sigmoid function, while for the categorical distributions in the constrained Poisson distribution, the piece-wise categorical distributions, as well as for the Gaussian-mixture model, the probabilities are given as logits with linear functions, which can be evaluated as probabilities using the softmax normalisation. Additionally for the piece-wise categorical distributions, we choose the cutoff *K* to be 4, since this strikes a good balance between the number of low and high counts for all examined data sets (see Supplementary Fig. S1 for an example of this). For the *k*-means clustering and the GMVAE model, the number of clusters is chosen to be equal to the number of classes, if cell types were provided.

The models are trained using one Monte Carlo sample for each example and using the Adam optimisation algorithm (Kingma and Ba, 2014) with a mini-batch size of 100 and a learning rate of 10^−4^. Additionally, we use batch normalisation (Ioffe and Szegedy, 2015) to improve convergence speed of the optimisation. We train all models for 500 epochs and use early stopping with the validation marginal log-likelihood lower bound to select parameters ***θ*** and ***ϕ***. We also use the warm-up optimisation scheme for 200 epochs for VAE and GMVAE models exhibiting overregularisation. To test whether the non-linearities in the neural network contributes to a better performance, we also run our models without hidden layers in the generative process. This corresponds to factor analysis (FA) with a generalised linear model link function. The optimal network architecture for the inference process is found in the same way as for the VAE models, and we similarly cluster the latent space using *k*-means clustering. In addition, we investigate the performance of a baseline linear (link) factor analysis (LFA). LFA assumes continuous data, so we first normalise the read counts using the counts-per-million method (Law *et al.*, 2014) and then apply log transformation to one plus the read count. We use a 25-dimensional Gaussian latent variable vector and cluster the latent space using *k*-means clustering. Both LFA and *k*-means clustering are performed using scikit-learn (Pedregosa *et al.*, 2011).

We compare our best-performing models with Seurat, scvis, and scVI for the PBMC data set. Seurat have in turn been compared to several other single-cell clustering methods (Duò *et al.*, 2018). Since Seurat uses the full data set, to make a fair comparison, we also train our best-performing models on the full data set. Because scvis is a visualisation method, we compare our models to it visually. In addition, we train our best-performing models using only two latent variables to visualise the latent space without using *t*-SNE. scVI is evaluated on the original counts similar to our method, so we can compare the test marginal log-likelihood lower bounds directly. We also perform *k*-means clustering on the latent space of scvis and scVI and compute the adjusted Rand indices of these clusterings to compare them with our method.

### 2.8 Software implementation

The models described above have been implemented in Python using the machine-learning software library TensorFlow (Abadi *et al.*, 2015) in an open-source and platform-independent software tool called scVAE (single-cell variational auto-encoders), and the source code is freely available online^5^ along with a user guide and examples. The RNA-seq data sets presented in Section 2.6 along with others are easily accessed using our tool. It also has extensive support for sparse data sets and can thus be used on the largest data sets currently available as demonstrated in Supplementary Section S4.

## 3 Results

Both the standard VAE and the GMVAE have been used to model both single-cell and bulk RNA-seq data sets (see Supplementary Sections S4–S5 for more). The performance of different likelihood functions have been investigated, and the latent spaces of the models have been examined.

### 3.1 scRNA-seq data of purified immune cells

We first modelled the scRNA-seq data set of gene expression counts for peripheral blood mononuclear cells (PBMC; Zheng *et al.*, 2017). To assess the optimal network architectures for these data sets, we first trained VAE-NB models with different network architecture as mentioned in Section 2.7 on the data set with all genes included. This was carried out as a grid search of number of hidden units and latent dimensions (Supplementary Fig. S2). We found that using two smaller hidden layers (of 100 units each) in the generative and inference processes yielded better results than only using one hidden layer, and a high-dimensional latent space of 100 dimensions resulted in the highest test marginal log-likelihood lower bound. A similar network architecture grid search were carried out using FA-NB models, and the optimal architecture was two large hidden networks of 1000 units each with a latent space of only 25 dimensions.

With these network architectures, different discrete likelihood functions were examined for the VAE, GMVAE, and FA models. The mean marginal log-likelihood lower bound and the mean adjusted Rand index for each model and likelihood function evaluated on the test set are plotted against each other in Figure 2 for a subset of model/link function combinations (see Supplementary Fig. S3 for the complete version). From this figure, it is clear that using a GM-VAE model with the negative binomial distribution yielded the highest lower bound as well as the highest Rand index, with the zero-inflated Poisson distribution as the next best. For each likelihood function, the GMVAE model yielded the highest lower bound, and in most cases, the distribution-based clustering of the GMVAE model outperforms *k*-means clustering in the latent space of the same model or of the other models on average. All scVAE models outperform the LFA baseline model. This shows that complex non-linear models really makes a difference for the marginal log-likelihood of the data. It should be noted, however, that there does not seem to be a strong positive correlation between Rand index and marginal likelihood lower bound. Similar conclusions are obtained using adjusted mutual information. The latent space of the GMVAE-NB model for the test set of the PBMC data set can be seen in Figure 3 (see Supplementary Fig. S4 for a larger version), which shows different clusters corresponding to distinct cell types, while more similar cell types are clustered close to each other or mixed together. In particular, cells cluster into well-known distinct immune cell populations: B cells, CD34+ cells, natural killer cells, and T cell sub-populations. Cytotoxic T cells, which express CD8, cluster together, and T cells expressing CD4 group together, while memory T cells, which can express both, are in a cluster between the two. Within the two CD4+ and CD8+ clusters, naïve and either regulatory or vanilla cells, respectively, form distinguishable groupings. Additionally, in the former cluster, helper T cells are mixed in with these two cell types, which makes sense, since their subtypes can express the same genes as either one.

**Fig. 2.**
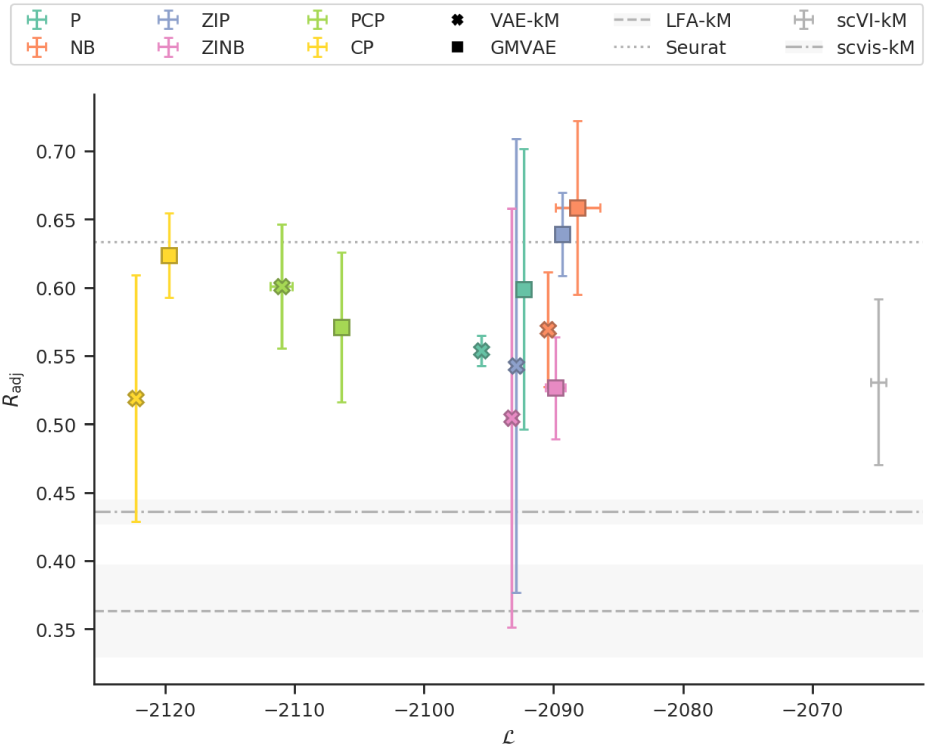
Comparison of VAE and GMVAE models with different likelihood functions trained and evaluated on the PBMC data set. For each combination the mean and the standard deviation of the adjusted Rand index are plotted against the marginal log-likelihood lower bound. These are compared to scVI and on the adjusted rand index only: LFA, Seurat (trained and evaluated on the full data set) and scvis.

**Fig. 3.**
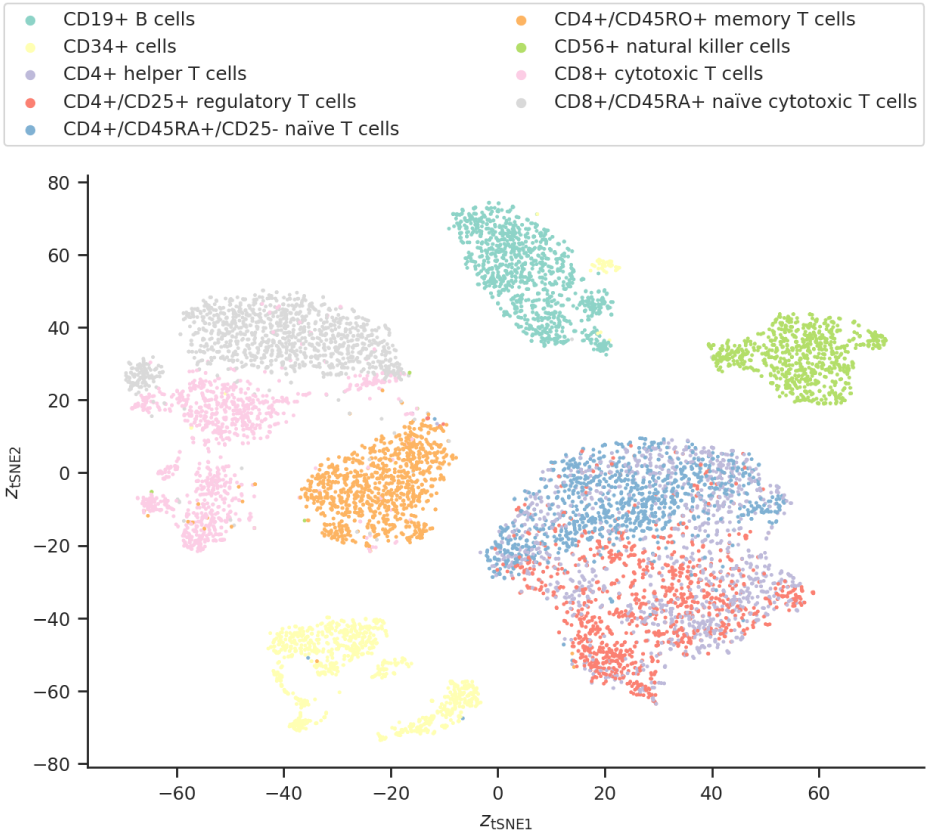
Latent space of the median-performing GMVAE-NB model trained and evaluated on the PBMC data set. The encoding of the cells in the latent space has been embedded in two dimensions using *t*-SNE and are colour-coded using their cell types. Clear separation corresponding to different cell types can be seen, but some similar cell types are also clustered close together or mixing together.

Furthermore, cells for several cell types seem to be split into subtypes, which could both be a biological and a batch effect. There is more overlap of the cell types in the latent spaces for the VAE-NB, FA-NB, and LFA models (Supplementary Fig. S5–7). It is also possible to visualise what each latent dimension encode about the cells by looking at how cell types are correlated with a given latent dimension (an example is shown in Supplementary Fig. S8). Finally, for the GMVAE model we investigated whether we could use the test marginal log-likelihood lower bound to identify the number of components. The result of scanning the number of components *K* from 5 to 13 is shown in Supplementary Fig. S9. The optimal value was found to be *K* = 9, which is the number of cell types in the data set.

The PBMC data set has also been analysed using Seurat at different resolutions (see Supplementary Table S1), and the highest adjusted Rand index obtained was *R*_adj_ = 0.634 at a resolution of 0.2. This yields nine clusters, which is also the number of cell types in the data set. This performance of Seurat have been included in Figure 2, and even though Seurat is trained on the full data set, two of the GMVAE models (using withheld data) perform better on average. A GMVAE-NB model was also trained on the full data set. This resulted in a lower bound of ℒ = −2046.0 ± 0.4 and a Rand index of *R*_adj_ = 0.656 ± 0.039 (see Supplementary Fig. S10 for its latent space), which is better than the best Rand index obtained using Seurat. In the latent space, we again find that the cells are clustered and arranged according to which CD molecule they express, and it also looks like some cells are clustered together in subtypes.

We also trained a scvis model with standard parameters as well as a GMVAE-NB model with a 2-d latent variable on the PBMC data set (see Supplementary Fig. S11 for their latent spaces). Comparing the latent space of the two models, both models succeed in separating out cells with dissimilar cell types, while struggling with cells of similar cell type, but the scvis model handles it better. This 2-d-latent-variable GMVAE-NB also has both a worse lower bound of ℒ= −2103.7 ±1.3 and a worse Rand index of *R*_adj_ = 0.420 ±0.014 compared to the GMVAE-NB using the optimal architecture, but comparable to the scvis, which has a Rand index of *R*_adj_ = 0.436 ±0.009 shown in Figure 2. The *t*-SNE of the latent space of this model also has a much better separation of the cells than the latent space of the scvis model.

Finally, we compared our method to scVI and the resulting mean lower bound and Rand index are shown in Figure 2. As can be seen from this figure, scVI significantly outperforms our models on the lower bound and are comparable to the standard VAE models in Rand index, but both comparable GMVAE models (with negative binomial distributions) are significantly better in ARI. Looking at the latent space of the scVI model (Supplementary Fig. S12), it is clear that the GMVAE model achieve more separation of the cells, especially of the T cells.

### 3.2 Bulk RNA-seq data of human cancer cells

The bulk RNA-seq data set of RSEM expected gene expression levels for human cancer cell samples from TCGA (Weinstein *et al.*, 2013) was also modelled. On a GPU with 12 GB of memory, we cannot model the data set using all genes with the optimal network architecture. Since the majority of the variance is captured in the 5000–10 000 most varying genes, we choose to limit the number of genes to 5000. As with the previous data set, different network architectures (Supplementary Fig. S13) and likelihood functions (Supplementary Fig. S14) were investigated for the VAE, GMVAE, and FA models. There is a clear difference in marginal log-likelihood lower bound between using either of the negative binomial distributions from any of the Poisson distributions, but for each likelihood function, the GMVAE model yielded the highest lower bound. *k*-means clustering on the VAE and GMVAE latent spaces for all likelihood functions resulted in the best adjusted Rand indices in general with the best one being for the GMVAE-P-kM model. Contrary to the sparse single cell data sets, for the TCGA data set only small performance gains were observed compared to the LFA baseline model. The latent space of the GMVAE-NB model for the test set of the TCGA data set (Supplementary Fig. S15) show that samples belonging to different tissue sites are clearly separated. The latent spaces of the VAE-NB, FA-NB, and LFA models look quite similar despite the difference in Rand index (Supplementary Fig. S16–18).

## 4 Discussion

We have shown that variational auto-encoders (VAEs) can be used to model single-cell and bulk RNA-seq data. The Gaussian-mixture VAE (GMVAE) model achieves higher marginal log-likelihood lower bounds on all data sets as well as a higher Rand indices for the PBMC data set compared to corresponding standard VAE, factor analysis (FA) and LFA models. The latent spaces for the GMVAE also showed the clearest separation according to cell type. Its built-in clustering helps the GMVAE achieve this.

In general we only find a weak correlation between test log marginal likelihood lower bound and Rand indices. Since likelihood is a measure of how well we are able to capture the multivariate transcript distribution and Rand index is a measure of alignment with predefined cell types the weak correlation simply implies that the are other important sources of variation in the data not directly related to cell type. Factor models (for count data and linear for transformed data) lead to significantly worse lower bounds, demonstrating that non-linear transformations can more easily express more subtle co-variation patterns in the data sets. We found that the negative binomial distribution modelled most data sets for most models the best, and that it was a close second to its zero-inflated version in the remaining cases. So explicit modelling of zero counts seems not be necessary. For much less sparse bulk RNA-seq dataset (TCGA), the advantage of working with the more advanced models were less pronounced. This might be explained by the diversity within each data sample and lesser need for accurately dealing with low and zero expression.

In modelling four different data sets, we have found the following guidelines help in achieving a higher marginal log-likelihood lower bound score on a data set. The network architecture should adapt to the data set: Use a large network when there are more cells and/or more cell types in the data set and when the data is less sparse. The models for all data sets also benefited from deeper network architectures. If the optimal network architecture is too large for computation, one can limit the number of genes to the most varying genes.^6^ For the likelihood function, try using the negative binomial distribution (or its zero-inflated version) first. In addition, the warm-up optimisation scheme proved valuable in avoiding overregularisation when modelling the scRNA-seq data sets.

Compared to Seurat (Satija *et al.*, 2015) the GMVAE-NB model achieved a higher adjusted Rand index on average for the PBMC data set. This was achieved working directly on the transcript counts with no preprocessing of the data. With variational auto-encoders being generative models, they can also model unseen data (test set or new data), whereas Seurat cannot. By modelling the cells with the GM-VAE model using the optimal configuration and then visualising the latent space using *t*-SNE, we also achieve a better visual representation of cells than the scvis model (Ding *et al.*, 2018). Even though the scVI model (Lopez *et al.*, 2018) achieves a better test marginal log-likelihood lower bound than any of our models on the PBMC data set, corresponding GM-VAE models still obtain latent spaces with more separate groupings. In Supplementary Section S4, we also showed that our method can scale up to very large data sets like the data set of 1.3 million mouse brain cells from 10x Genomics.

## 5 Summary and conclusion

We show that two variations of the variational auto-encoder are able to model gene expression counts using appropriate discrete probability distribution as likelihood (link) functions and provide a software implementation. These models are probabilistic and put a lower bound on the marginal likelihood of the data sets, enabling us to perform likelihood-based model comparison. We have applied both models successfully to single-cell and – to a lesser degree – bulk RNA-seq data sets, and the GMVAE model achieves better clustering and representation of the cells than Seurat, scvis, scVI, and a baseline method using size-factor normalisation and log transformation, linear factor analysis, and *k*-means clustering. However, scVI achieves a better bound for the likelihood, which might be attributed to including the sequencing depth as a latent variable in the scVI model. Building clustering into the Gaussian-mixture variational auto-encoder, we have a model that can cluster cells into cell populations without the need of a pipeline with several parameter selection steps. Having both an inference process and a generative process, makes it possible to project new data onto an existing latent space, or even simulate new data from samples in the latent space. This means that new cells can be introduced to an already trained model, and it could enable combining the latent representations of two cells to generate a cell and the transitional states in-between. In the scVI model, a zero-inflated negative binomial distribution is used. We observe that using a negative binomial distribution overall is a better model for our method. As in Lopez *et al.* (2018), we also provide batch correction within our framework.

As for future extensions we would also like to make the models more flexible by adding more latent variable layers (Sønderby *et al.*, 2016) and make the models learn the number of clusters (Rasmussen, 2000) rather than setting it a priori. We could also use semi-supervised learning and active learning to better classify cells and identify cell populations and associated response variables. This would also help with transfer learning enabling modelling multiple data sets with the same model. As mentioned above, it should also be possible to combine encodings of the cells in the latent space and produce in-between cells like Lotfollahi *et al.* (2018). We would also like to extent our investigation of what dimensions of the latent variables encode (Kinalis *et al.*, 2019). We note that it is possible to apply these models to data sets with multiple modalities such as RNA-seq and exome sequencing (Brouwer and Lió, 2017).

## Supporting information

Supplementary matierals

## Acknowledgements

This work was supported by the Lundbeck Foundation to THP and the Novo Nordisk Foundation to THP. We would like to thank Lars Maaløe for discussions about multimodal VAEs and technical help for implementing these. We also want to thank Aki Vehtari for discussions about discrete probability distributions and model comparison. Lastly, we gratefully acknowledge the support of NVIDIA Corporation with the donation of Tesla Titan Xp GPUs used for this research.

## Footnotes

1 A list of software packages for single-cell data analysis is available at https://github.com/seandavi/awesome-single-cell.

2 Detailed in a blog post by Rui Shu: http://ruishu.io/2016/12/25/gmvae/.

3 Available online at https://support.10xgenomics.com/single-cell-gene-expression/datasets/ (under “Single Cell 3′ Paper: Zheng et al. 2017”).

4 Available online at https://xenabrowser.net/datapages/?dataset=tcga_gene_expected_count&host=https://toil.xenahubs.net.

5 https://github.com/scvae/scvae.

6 We did this for the bulk RNA-seq data set, and the optimal network architecture stayed the same.

