## Supplementary matierals for "scVAE: Variational auto-encoders for single-cell gene expression data"

#### SUPPLEMENTARY MATERIALS

**Christopher Heje Grønbech, Maximillian Fornitz Vording, Pascal Timshel, Casper Kaae Sønderby, Tune Hannes Pers, and Ole Winther**

#### S1 Inference for the Gaussian-mixture VAE

To compute the marginal likelihood for the GMVAE, we must sum and average out the latent variables, $y$ and $\mathbf{z}$:

$$p_{\boldsymbol{\theta}}(\mathbf{x}) = \sum_{y} \int p_{\boldsymbol{\theta}}(\mathbf{x}, y, \mathbf{z}) \, \mathrm{d}\mathbf{z} \tag{S1}$$

with $p_{\boldsymbol{\theta}}(\mathbf{x}, y, \mathbf{z}) = p_{\boldsymbol{\theta}}(\mathbf{x}|\mathbf{z}) p_{\boldsymbol{\theta}}(\mathbf{z}|y) p_{\boldsymbol{\theta}}(y)$. Because of the additional latent variable, the marginal log-likelihood lower bound for the GMVAE has an additional marginalisation of the the discrete categorical random variable $y \in \{1, \ldots, K\}$. We use the following variational distribution analogously to their corresponding prior distributions in Equations (6) in the main text:

$$q_{\boldsymbol{\phi}}(\mathbf{z}, y|\mathbf{x}) = q_{\boldsymbol{\phi}}(\mathbf{z}|\mathbf{x}, y) \, q_{\boldsymbol{\phi}}(y|\mathbf{x}), \tag{S2a}$$

$$q_{\boldsymbol{\phi}}(\mathbf{z}|\mathbf{x}, y) = \mathcal{N}(\mathbf{z}; \boldsymbol{\mu}_{\boldsymbol{\phi}}(\mathbf{x}, y), \boldsymbol{\sigma}^2_{\boldsymbol{\phi}}(\mathbf{x}, y)\mathbf{I}), \tag{S2b}$$

$$q_{\boldsymbol{\phi}}(y|\mathbf{x}) = \mathrm{Cat}(y; \boldsymbol{\pi}_{\boldsymbol{\phi}}(\mathbf{x})). \tag{S2c}$$

Here, $\boldsymbol{\mu}_{\boldsymbol{\phi}}(\mathbf{x}, y)$, $\boldsymbol{\sigma}^2_{\boldsymbol{\phi}}(\mathbf{x}, y)$, $\boldsymbol{\pi}_{\boldsymbol{\phi}}(\mathbf{x})$ are non-linear transformations of $\mathbf{x}$ and $y$ parameterised by $\boldsymbol{\theta}$. For example, using a single-layer feed-forward neural network for each, they are computed as

$$\boldsymbol{\mu}_{\boldsymbol{\phi}}(\mathbf{x}, y) = \mathbf{u}\Big(\mathbf{W}^{(\boldsymbol{\phi})}_{\boldsymbol{\mu},y}\mathbf{x} + \mathbf{b}^{(\boldsymbol{\phi})}_{\boldsymbol{\mu},y}\Big), \tag{S3a}$$

$$\boldsymbol{\sigma}^2_{\boldsymbol{\phi}}(\mathbf{x}, y) = \mathbf{v}\Big(\mathbf{W}^{(\boldsymbol{\phi})}_{\boldsymbol{\sigma}^2,y}\mathbf{x} + \mathbf{b}^{(\boldsymbol{\phi})}_{\boldsymbol{\sigma}^2,y}\Big), \tag{S3b}$$

$$\boldsymbol{\pi}_{\boldsymbol{\phi}}(\mathbf{x}) = \mathbf{w}\Big(\mathbf{W}^{(\boldsymbol{\phi})}_{\boldsymbol{\pi}}\mathbf{x} + \mathbf{b}^{(\boldsymbol{\phi})}_{\boldsymbol{\pi}}\Big), \tag{S3c}$$

where $\mathbf{u}(\cdot)$, $\mathbf{v}(\cdot)$, $\mathbf{w}(\cdot)$ are appropriate non-linear transformations. As for $p(y)$, $\boldsymbol{\pi}_{\boldsymbol{\phi}}(\mathbf{x})$ is $K$-dimensional probability vector, and its $k$th component, $\pi^{(\boldsymbol{\theta})}_k(\mathbf{x})$, is the responsibility of the $k$th Gaussian distribution for $\mathbf{x}$, quantifying how much this distribution contributes to the overall probability of $\mathbf{x}$.

The variational bound becomes:

$$\begin{aligned}\mathcal{L}(\boldsymbol{\theta}, \boldsymbol{\phi}; \mathbf{x}) = \mathrm{E}_{q_{\boldsymbol{\phi}}(y|\mathbf{x})}\big[&\mathrm{E}_{q_{\boldsymbol{\phi}}(\mathbf{z}|\mathbf{x},y)}[\log p_{\boldsymbol{\theta}}(\mathbf{x}|\mathbf{z})] - \mathrm{KL}(q_{\boldsymbol{\phi}}(\mathbf{z}|\mathbf{x}, y) \,\|\, p_{\boldsymbol{\theta}}(\mathbf{z}|y))\big] \\ &- \mathrm{KL}(q_{\boldsymbol{\phi}}(y|\mathbf{x}) \,\|\, p(y)).\end{aligned} \tag{S4}$$

The first term is the standard VAE lower bound for each class $y$ averaged over all classes, and the second term is the KL divergence for $y$. The approximate posterior distributions in the inference process are formulated The inference process for the GMVAE is shown in Fig. 1B of the main text,

The marginal log-likelihood lower bound can further be modified to:

$$\begin{aligned}
\mathcal{L}(\boldsymbol{\theta}, \boldsymbol{\phi}; \mathbf{x}) &= \mathrm{E}_{q_{\boldsymbol{\phi}}(y|\mathbf{x})}\left[\mathrm{E}_{q_{\boldsymbol{\phi}}(\mathbf{z}|\mathbf{x},y)}[\log p_{\boldsymbol{\theta}}(\mathbf{x}|\mathbf{z})] - \mathrm{KL}(q_{\boldsymbol{\phi}}(\mathbf{z}|\mathbf{x},y) \,\|\, p_{\boldsymbol{\theta}}(\mathbf{z}|y))\right] \\
&\quad - \mathrm{KL}(q_{\boldsymbol{\phi}}(y|\mathbf{x}) \,\|\, p(y)) \\
&= \sum_{k=1}^{K} q_{\boldsymbol{\phi}}(y=k|\mathbf{x})\big(\mathrm{E}_{q_{\boldsymbol{\phi}}(\mathbf{z}|\mathbf{x},y=k)}[\log p_{\boldsymbol{\theta}}(\mathbf{x}|\mathbf{z})] - \mathrm{KL}(q_{\boldsymbol{\phi}}(\mathbf{z}|\mathbf{x},y=k) \,\|\, p_{\boldsymbol{\theta}}(\mathbf{z}|y=k))\big) \\
&\quad - \sum_{k=1}^{K} q_{\boldsymbol{\phi}}(y=k|\mathbf{x}) \log \frac{q_{\boldsymbol{\phi}}(y=k|\mathbf{x})}{p_{\boldsymbol{\theta}}(y=k)} \\
&= \sum_{k=1}^{K} \pi_k^{(\boldsymbol{\theta})}(\mathbf{x})\big(\mathrm{E}_{q_{\boldsymbol{\phi}}(\mathbf{z}|\mathbf{x},y=k)}[\log p_{\boldsymbol{\theta}}(\mathbf{x}|\mathbf{z})] - \mathrm{KL}(q_{\boldsymbol{\phi}}(\mathbf{z}|\mathbf{x},y=k) \,\|\, p_{\boldsymbol{\theta}}(\mathbf{z}|y=k))\big) \\
&\quad - \sum_{k=1}^{K} \pi_k^{(\boldsymbol{\theta})}(\mathbf{x}) \log \frac{\pi_k^{(\boldsymbol{\theta})}(\mathbf{x})}{\pi_k}.
\end{aligned} \tag{S5}$$

The reconstruction $\tilde{\mathbf{x}}$ is computed as: $\tilde{\mathbf{x}} = \int \sum_y \sum_{\mathbf{x}'} \mathbf{x}' p(\mathbf{x}' \mid \mathbf{z}) q(\mathbf{z} \mid \mathbf{x}, y) q(y \mid \mathbf{x}) d\mathbf{z}$.

### S2 Avoiding overregularisation

Maximising the marginal log-likelihood lower bound for the VAE model in Equation (11) of the main text is a compromise between either maximising the reconstruction error or minimising the KL divergence. This leads the model to balance reconstructing the signal exactly and learning a latent representation, which follows the prior distribution completely. As such, the KL divergence regularises the model, thereby making the model more generalisable to unseen data. But during the initial training phase, the regularisation can be too strong, constricting the model to learn for only a few of the components of the latent variable (Bowman *et al.*, 2016; Sønderby *et al.*, 2016). This is undesirable, since the latent variable can then only use a small subset of its components to represent the input data. Following Bowman *et al.* (2016) and Sønderby *et al.* (2016), we introduce a weight $\beta$ for the KL divergence to alleviate this:

$$\mathcal{L}_W(\boldsymbol{\theta}, \boldsymbol{\phi}; \mathbf{x}) = \mathrm{E}_{q_{\boldsymbol{\phi}}(\mathbf{z}|\mathbf{x})}[\log p_{\boldsymbol{\theta}}(\mathbf{x}|\mathbf{z})] - \beta\, \mathrm{KL}(q_{\boldsymbol{\phi}}(\mathbf{z}|\mathbf{x}) \,\|\, p_{\boldsymbol{\theta}}(\mathbf{z})). \tag{S6}$$

$\beta$ is gradually increased linearly from 0 to 1 during the initial $W$ epochs of training. Thus, the model goes from a traditional auto-encoder to a full variational auto-encoder after $W$ epochs. This optimisation scheme is referred to as warm-up by Sønderby *et al.* (2016).

### S3 Per-dimension KL divergence

When we inspect the latent representations of the model to derive a biological interpretation, we would like to have a way to order them. For the GMVAE model, latent dimensions where the posterior latent distribution matches the Gaussian mixture prior well are good candidates for this, because these will have a bigger tendency to cluster the data. To derive a per-dimension expression, we can start by defining the KL for the $y$th component:

$$\mathrm{KL}(q_{\boldsymbol{\phi}}(\mathbf{z}|\mathbf{x},y) \,\|\, p_{\boldsymbol{\theta}}(\mathbf{z}|y)). \tag{S7}$$

We can further average this over the component posterior:

$$\mathrm{E}_{q_{\boldsymbol{\phi}}(y|\mathbf{x})}[\mathrm{KL}(q_{\boldsymbol{\phi}}(\mathbf{z}|\mathbf{x},y) \,\|\, p_{\boldsymbol{\theta}}(\mathbf{z}|y))]. \tag{S8}$$

This expression will emphasise the KL divergence of components with high responsibility for the data point. Next we can use that the distributions over $\mathbf{z}$ are conditional independent, so we can write this as a sum over components:

$$\sum_l \mathrm{E}_{q_{\boldsymbol{\phi}}(y|\mathbf{x})}[\mathrm{KL}(q_{\boldsymbol{\phi}}(z_l|\mathbf{x},y) \,\|\, p_{\boldsymbol{\theta}}(z_l|y))] \tag{S9}$$

and define

$$C_l \equiv \mathrm{E}_{q_{\boldsymbol{\phi}}(y|\mathbf{x})}[\mathrm{KL}(q_{\boldsymbol{\phi}}(z_l|\mathbf{x},y) \,\|\, p_{\boldsymbol{\theta}}(z_l|y))] \tag{S10}$$

as the per-dimension averaged KL divergence. The $R$-samples expression for $C_l$ can be written as:

$$C_l = \sum_{k=1}^{K} q_{\boldsymbol{\phi}}(y = k|\mathbf{x}) \sum_{r=1}^{R} \log \frac{q_{\boldsymbol{\phi}}(z_{rl}|\mathbf{x})}{p_{\boldsymbol{\theta}}(z_{rl}|y = k)}. \tag{S11}$$

We use this expression averaged over the data set to sort latent dimensions in ascending order.

### S4 scRNA-seq data of mouse brain cells

Another single-cell gene expression data set made publicly available by 10x Genomics[1] is also modelled, and as for the PBMC data set (Section 3.1 in the main text), we use the filtered gene–cell matrix. This data set contains gene expression levels for 1.3 million mouse brain cells (MBC), but in contrast to the PBMC data set, the cells were not purified, so this data set does not contain any information about cell types. A smaller subset of 20 000 uniformly randomly selected cells (MBC-20k) is also provided by 10x Genomics. Both data sets are summarised in Table S2.

#### S4.1 Results

We can scale our method to train on the full MBC data set without any modifications. However, because of the large number of cells, we used the smaller subset of 20 000 uniformly randomly selected cells to perform a network architecture grid search (Fig. S19) as well as test the different likelihood functions and models (Fig. S20) following the same procedure as for other data sets. Since no cell types were available for this data set, neither the adjusted Rand index nor adjusted mutual information could be evaluated. For each likelihood function, the GMVAE resulted in the highest test marginal log-likelihood lower bound, and the highest among these were achieved using the negative binomial distribution with its zero-inflated version as a close second. The latent space of the GMVAE-NB model for the test set of the MBC-20k data set can be seen in Fig. S21.

For the full data set, a GMVAE model were trained using the optimal architecture found for the subset and zero-inflated negative binomial distribution, and its latent space is plotted in in Fig. S22, which shows somewhat clear regions for each cluster and even some separation for some clusters. Training the GMVAE model one epoch on the full data set took approximately 26 minutes on a GeForce GTX 1080 Ti graphics card.

### S5 Bulk RNA-seq data of tissue-specific human cells

We also modelled a bulk RNA-seq data set of gene expression count for human cells from different tissue sites made publicly available by Genotype-Tissue Expression (GTEx) Consortium[2] (Lonsdale *et al.*, 2013). It is summarised in Table S2. This data set contains gene expression counts for samples of human cells from 51 tissue sites and two cell lines. We collate cell samples from similar tissue site, and end up with 31 cell populations, which we use when plotting and for computing the Rand index. Gene IDs are used in this data set, so the available mapping from the GTEx Consortium is used to map the IDs to gene names. The difference in number of genes and sparsity from the original data set is not large.

#### S5.1 Results

Different network architectures (Fig. S23) as well as likelihood functions and models (Fig. S24) were also investigated for this data set. Because of same memory limitations as for the TCGA data set (see Section 3.2 in the main text), we limited the number of genes to the 5000 most varying genes. The results are also similar to those for the TCGA data set with four noticeable exceptions: (a) It is the GMVAE-ZINB model which has the highest marginal log-likelihood lower bound closely followed by the GMVAE-NB model; (b) the highest adjusted Rand index is achieved by the GMVAE-km-PCP model; (c) the constrained Poisson distribution is able to model this data set, and the CP models are close in lower-bound performance to the NB and ZINB models relative to the remaining Poisson models, while they achieve similar Rand indices to the NB and ZINB models; and (d) $k$-means

[1] Available online at `https://support.10xgenomics.com/single-cell-gene-expression/datasets/1.3.0/1M_neurons`.

[2] Available online at `https://gtexportal.org/`.

clustering is marginally better on average than the LFA baseline model for every model. The latent space of the GMVAE-ZINB model for the test set of the GTEx data set can be seen in Fig. S25.

### S6 Supplementary Tables

**Table S1.** Number of clusters found, $C_{\mathsf{est}}$, and adjusted Rand index, $R_{\mathsf{adj}}$ for the PBMC data set using Seurat with different resolutions (the best results have been highlighted in bold).

| Resolution | $C_{\text{est}}$ | $R_{\text{adj}}$ |
|---|---|---|
| 0.1 | 7 | 0.577 784 |
| 0.2 | **9** | **0.633 611** |
| 0.3 | 10 | 0.631 494 |
| 0.4 | 10 | 0.595 873 |
| 0.5 | 12 | 0.592 172 |
| 0.6 | 15 | 0.508 233 |
| 0.7 | 15 | 0.509 282 |
| 0.8 | 16 | 0.542 031 |
| 1 | 18 | 0.488 330 |
| 2 | 32 | 0.451 832 |
| 4 | 45 | 0.431 569 |
| 8 | 84 | 0.515 057 |

**Table S2.** Overview of gene expression data sets.

| Data set | Examples | Features | Classes | Sparsity [%] |
|---|---|---|---|---|
| PBMC | 92 043 | 32 738 | 9[a] | 98.05 |
| MBC | 1 306 127 | 27 998 | — | 92.82 |
| MBC-20k | 20 000 | 27 998 | — | 92.82 |
| TCGA | 10 830 | 58 581 | 29 | 52.47 |
| GTEx | 11 688 | 54 271 | 53 | 47.16 |

[a] CD19+ B cells, CD34+ cells, CD4+ helper T cells, CD4+/CD25+ regulatory T cells, CD4+/CD45RA+/CD25- naïve T cells, CD4+/CD45RO+ memory T cells, CD56+ natural killer cells, CD8+ cytotoxic T cells, CD8+/CD45RA+ naïve cytotoxic T cells.

#### S7 Supplementary Figures

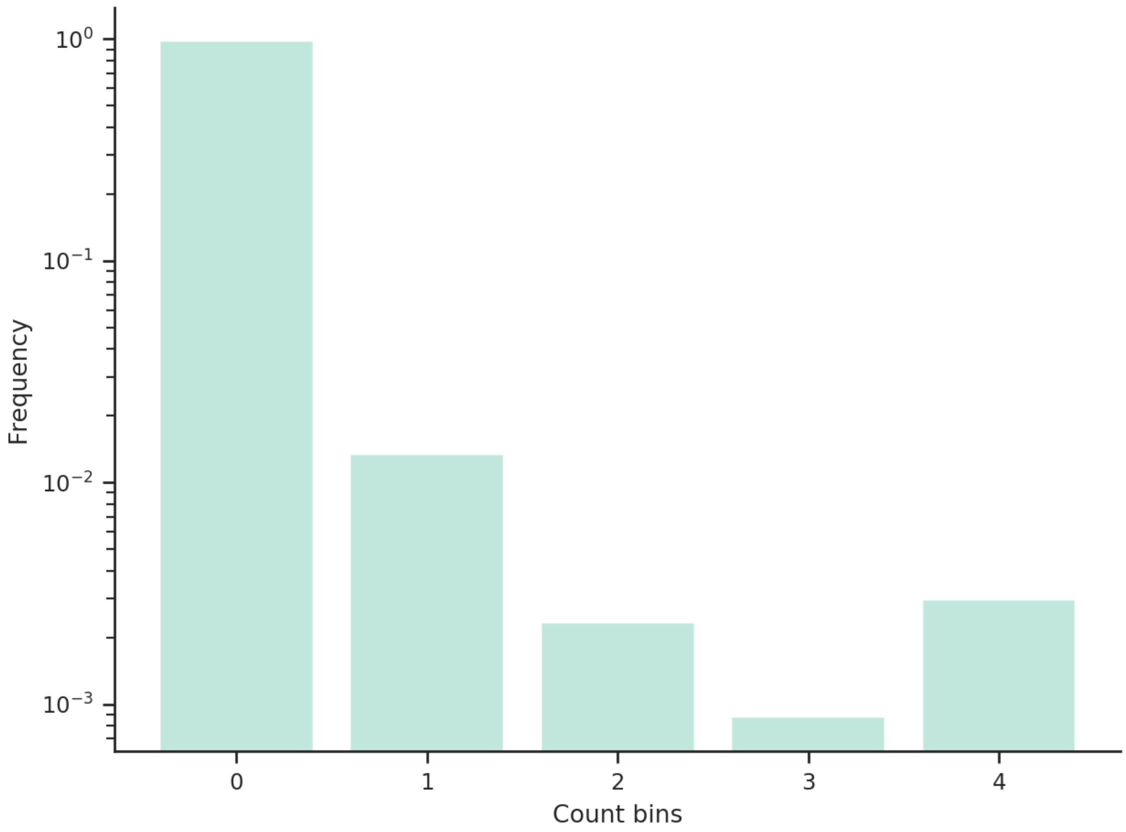


**Figure S1.** Gene expression count histogram for the PBMC data set. Counts above $K = 4$ having been set to this value.

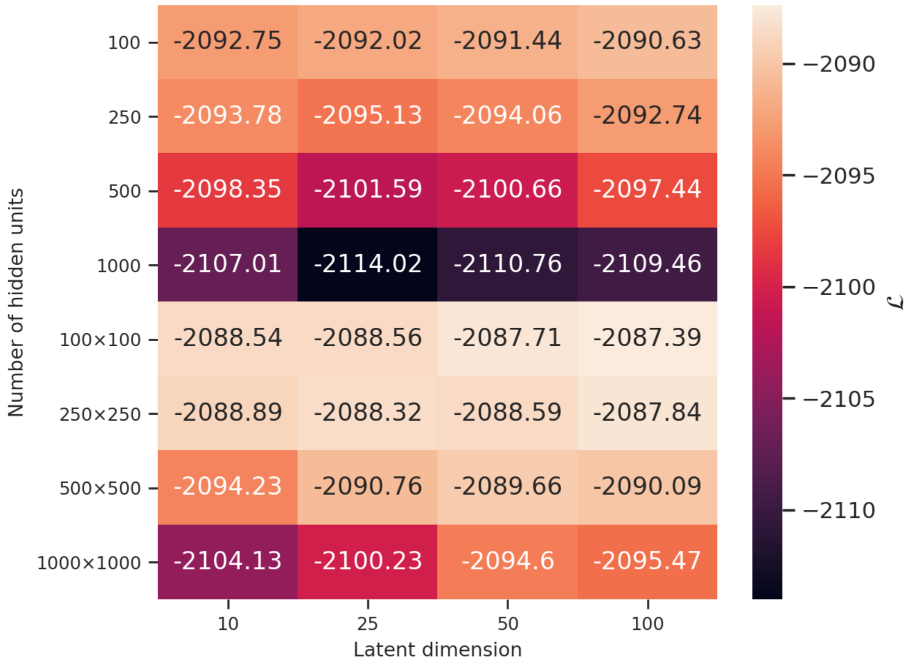


**Figure S2.** Test marginal log-likelihood lower bounds, $\mathcal{L}$, for the PBMC data set using a VAE-NB model with different latent dimensions and different number of hidden units. The multiplication sign ($\times$) separate the number of units in each hidden layer. Two hidden layers of 100 units with 100 latent dimensions are the most optimal.

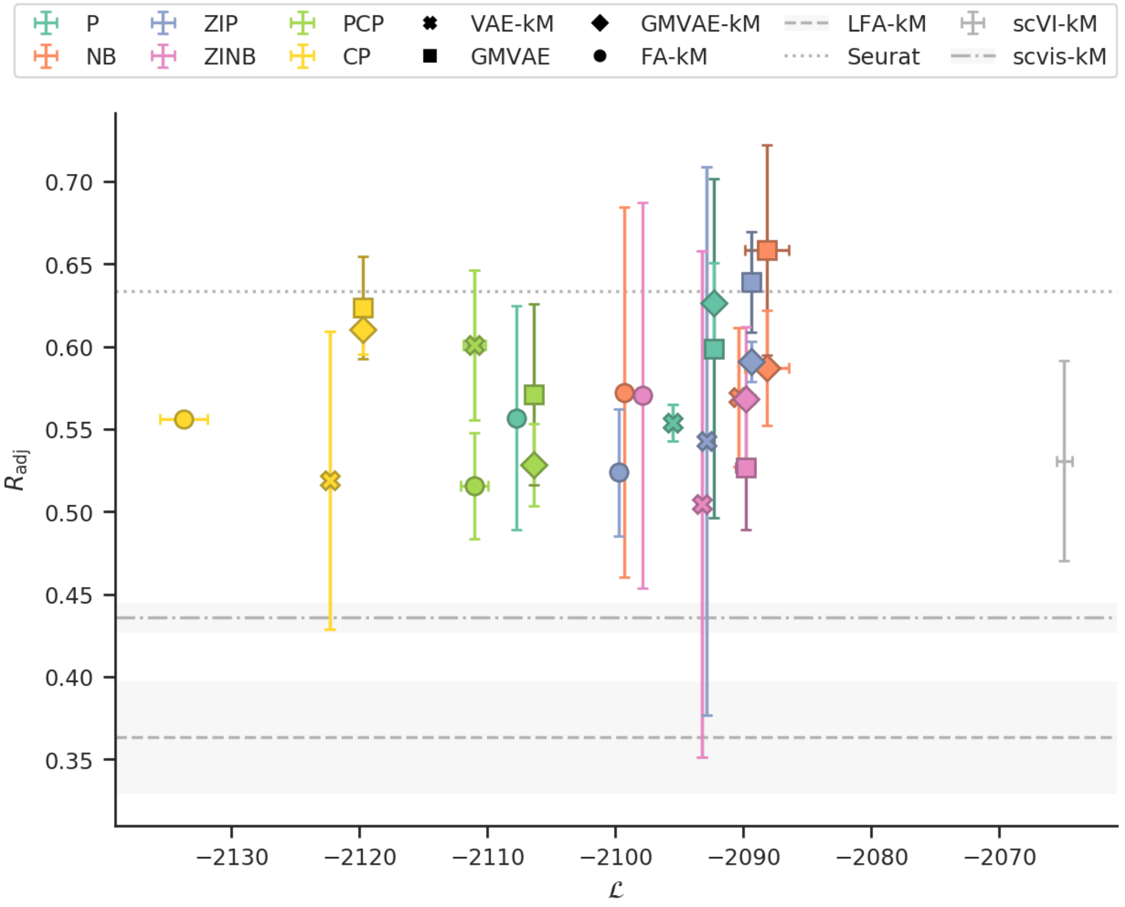


**Figure S3.** Comparison of VAE, GMVAE, and FA models with different likelihood functions trained and evaluated on the PBMC data set. For each combination the mean and the standard deviation of the adjusted Rand index is plotted against the marginal log-likelihood lower bound. The mean and the standard deviation of the adjusted Rand index is also plotted for the LFA baseline model as well as for the Seurat (trained and evaluated on the full data set), scvis, and scVI (against the marginal log-likelihood lower bound) methods.

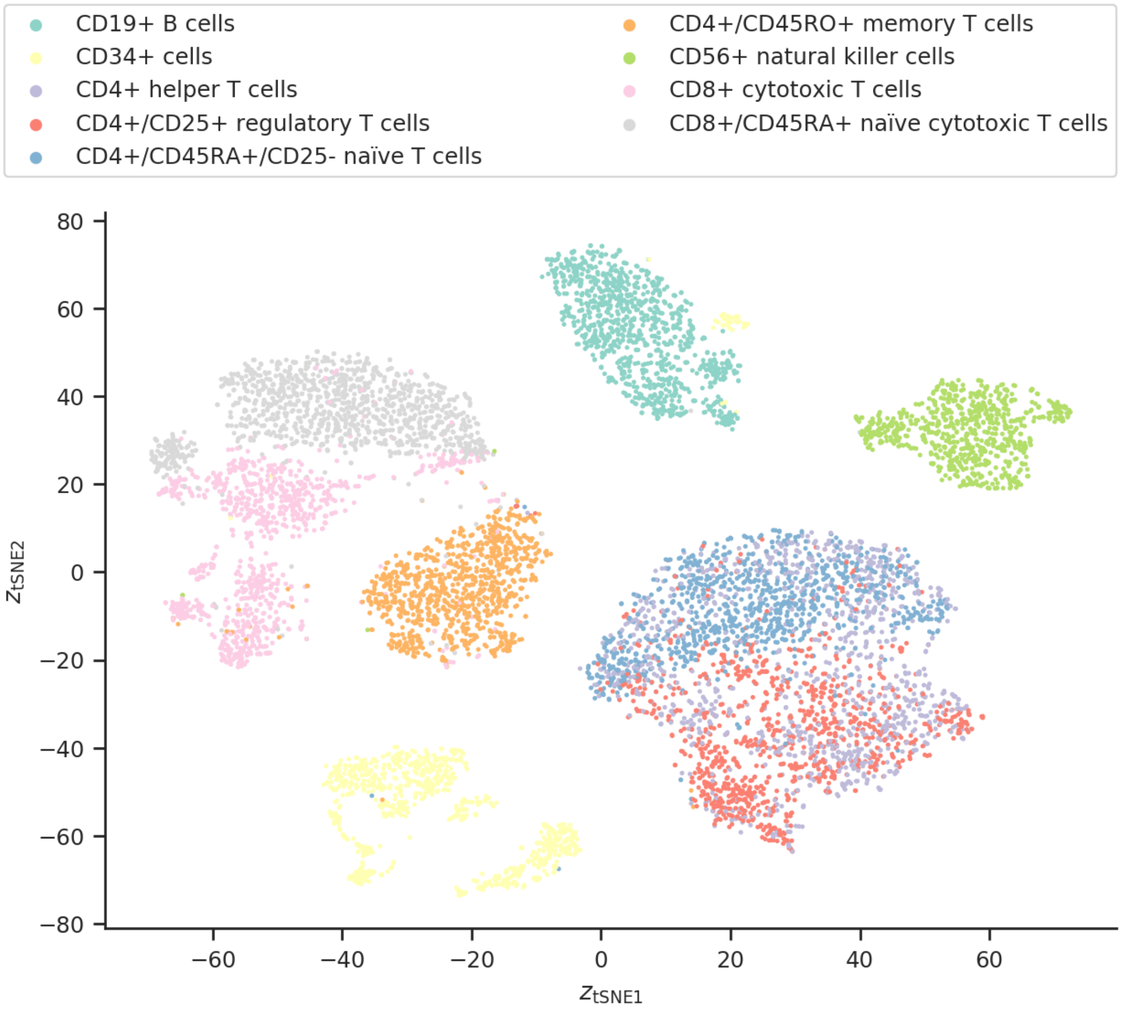


**Figure S4.** Latent space of the median-performing GMVAE-NB model trained and evaluated on the PBMC data set. The encoding of the cells in the latent space has been embedded in two dimensions using *t*-SNE and are colour-coded using their cell types. Clear separations can be seen corresponding to different cell types, but some similar cell types are also clustered close together or mixing together.

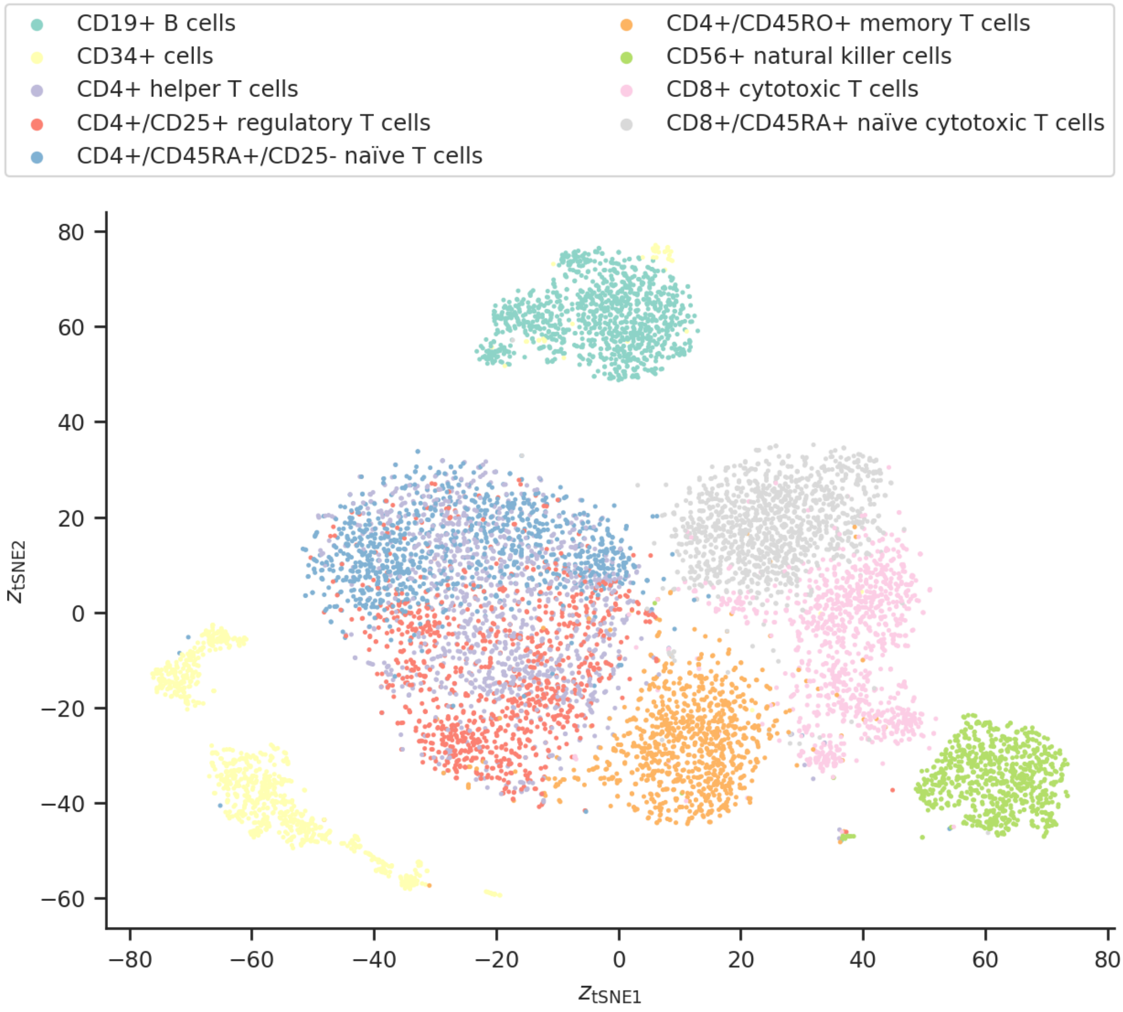


**Figure S5.** Latent space of the median-performing VAE-NB model trained and evaluated on the PBMC data set. The encoding of the cells in the latent space has been embedded in two dimensions using *t*-SNE and are colour-coded using their cell types. Some separations can be seen corresponding to different cell types, and some similar cell types are also clustered close together or mixing together. However, there is more overlap than for the GMVAE model.

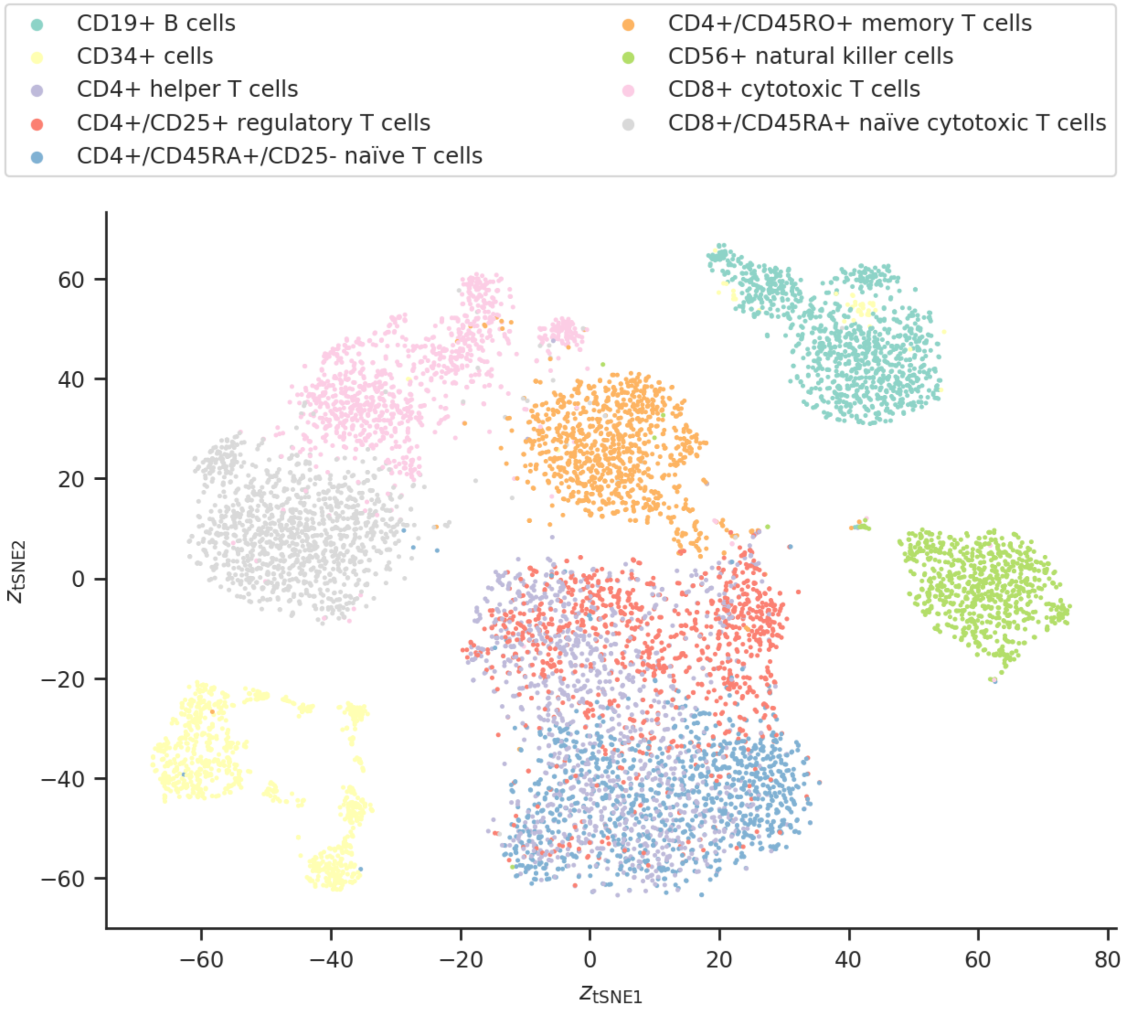


**Figure S6.** Latent space of the median-performing FA-ZINB model trained and evaluated on the PBMC data set. The encoding of the cells in the latent space has been embedded in two dimensions using *t*-SNE and are colour-coded using their cell types. Some separations can be seen corresponding to different cell types, and some similar cell types are also clustered close together or mixing together. However, there is more overlap than for the GMVAE model.

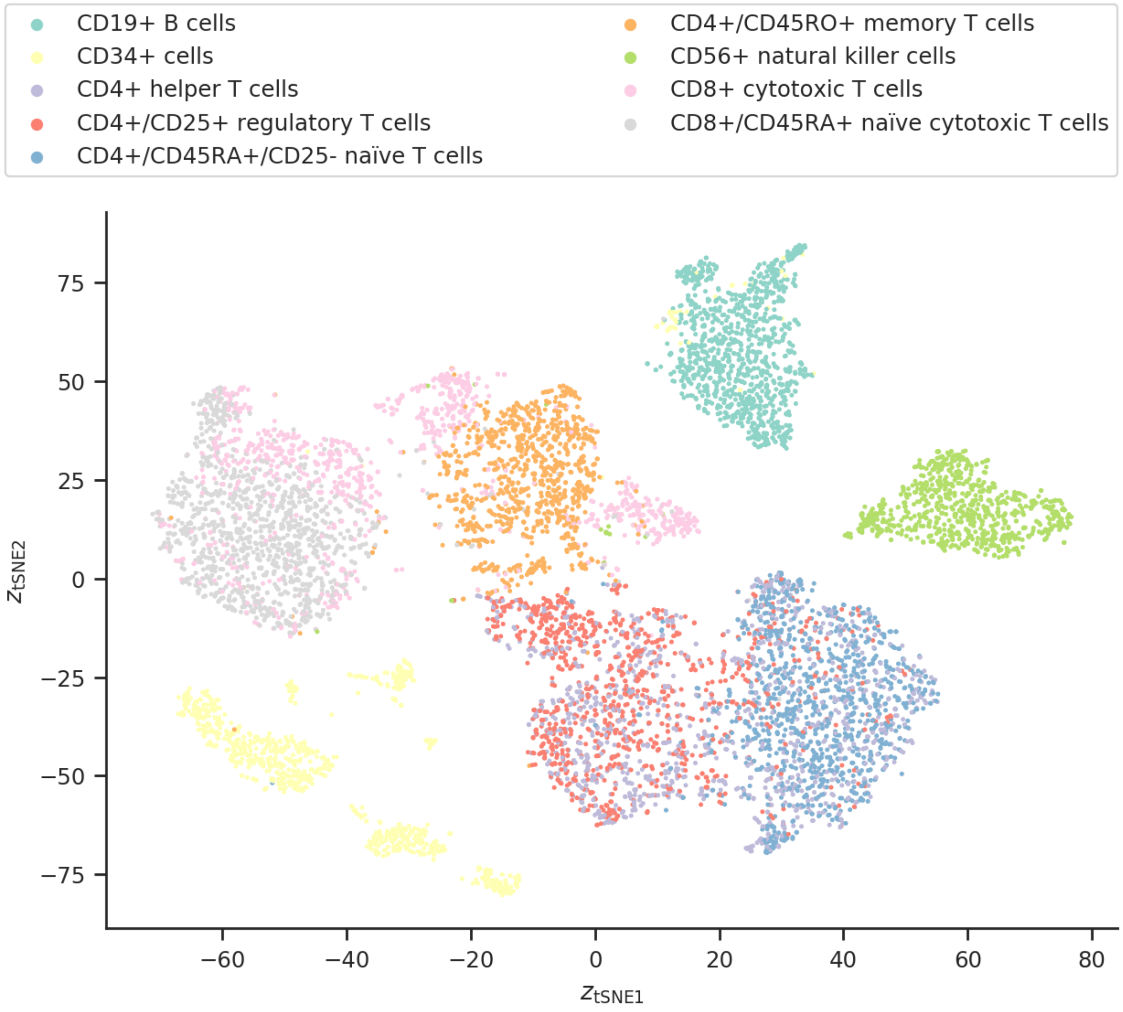


**Figure S7.** Latent space of the median-performing LFA baseline model trained and evaluated on the PBMC data set. The encoding of the cells in the latent space has been embedded in two dimensions using *t*-SNE and are colour-coded using their cell types. Some separations can be seen corresponding to different cell types, and some similar cell types are also clustered close together or mixing together. However, there is more overlap than for the GMVAE model.

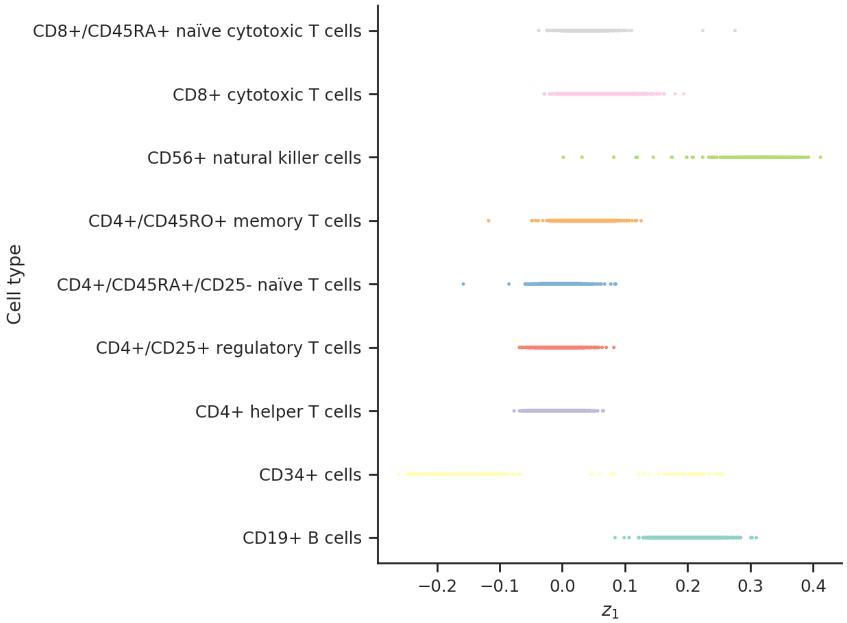


**(a)**

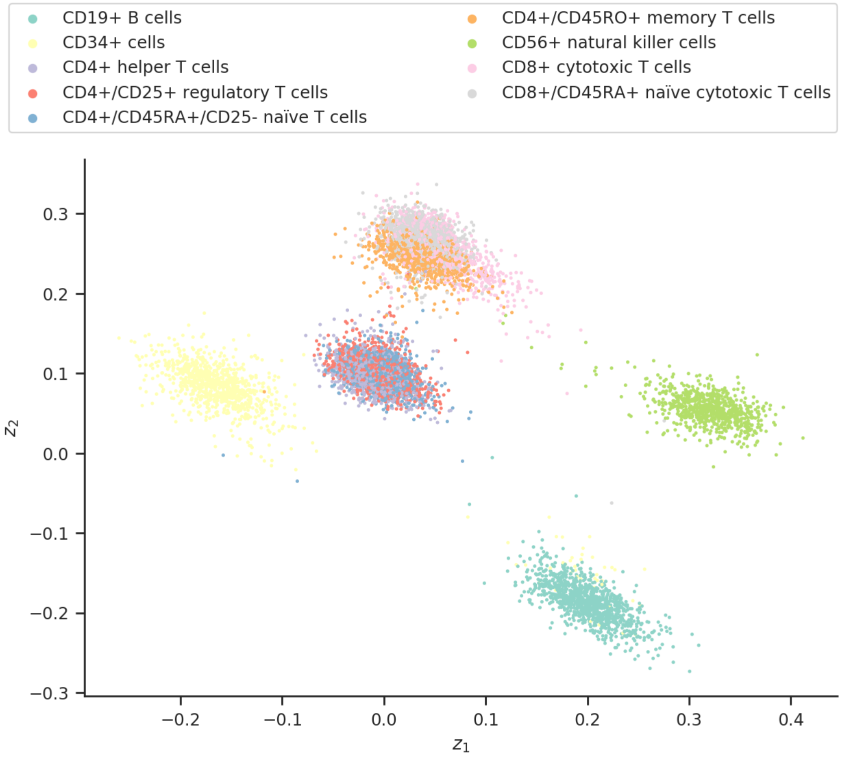


**(b)**

**Figure S8.** Scatter plots between **(a)** cell types and the dimension of the latent variable of $\mathbf{z}$ as well as **(b)** the two first dimensions of the latent variable $\mathbf{z}$. The dimensions of the latent variable $\mathbf{z}$ are sorted by their KL divergence, so the first dimensions have the lowest KL divergence, meaning that they deviate the least between the prior and posterior distributions for $\mathbf{z}$ (see Section S3). The latter plot shows some of the same clusters found in Figure S4, but only using two dimensions of $\mathbf{z}$. The encoding of the cells in the latent space has been embedded in two dimensions using $t$-SNE and are colour-coded using their cell types.

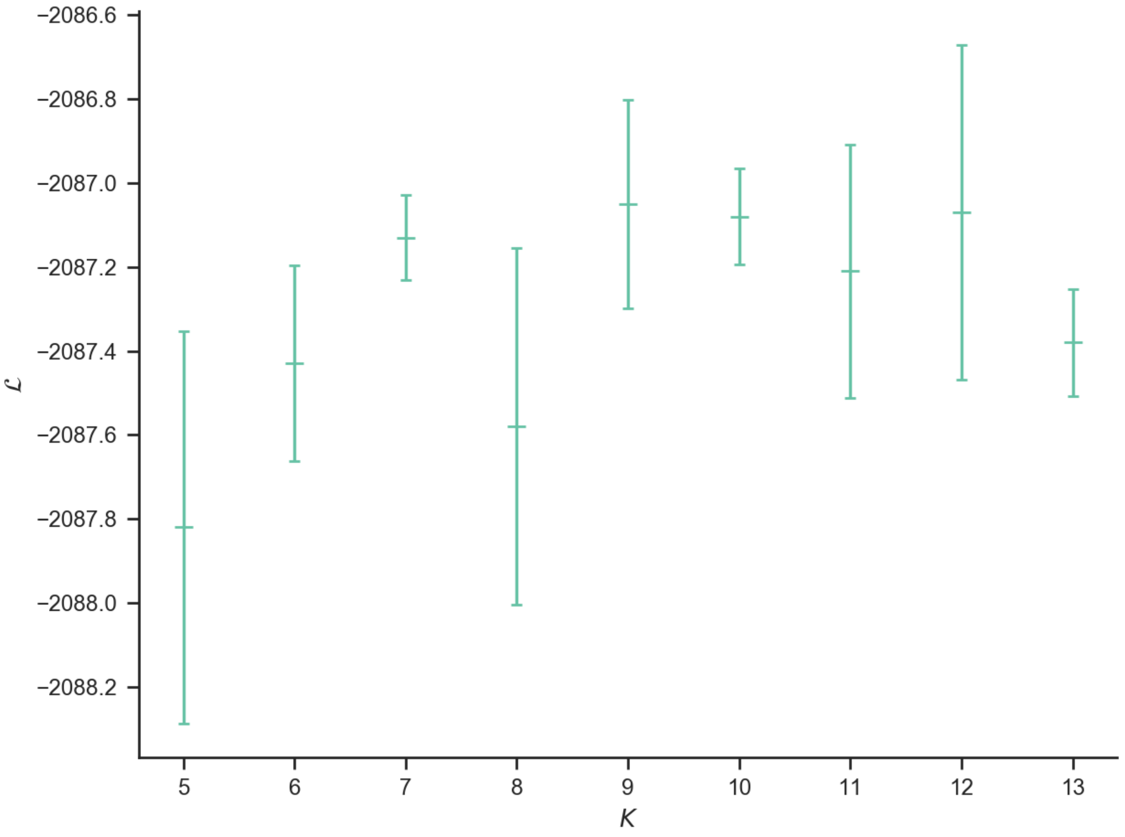


**Figure S9.** Comparison of test marginal log-likelihood lower bounds, $\mathcal{L}$, for the GMVAE-NB-model trained and evaluated on the PBMC data set for different number of components, $K$. The optimal value was found to be $K = 9$, which is the number of cell types in the data set.

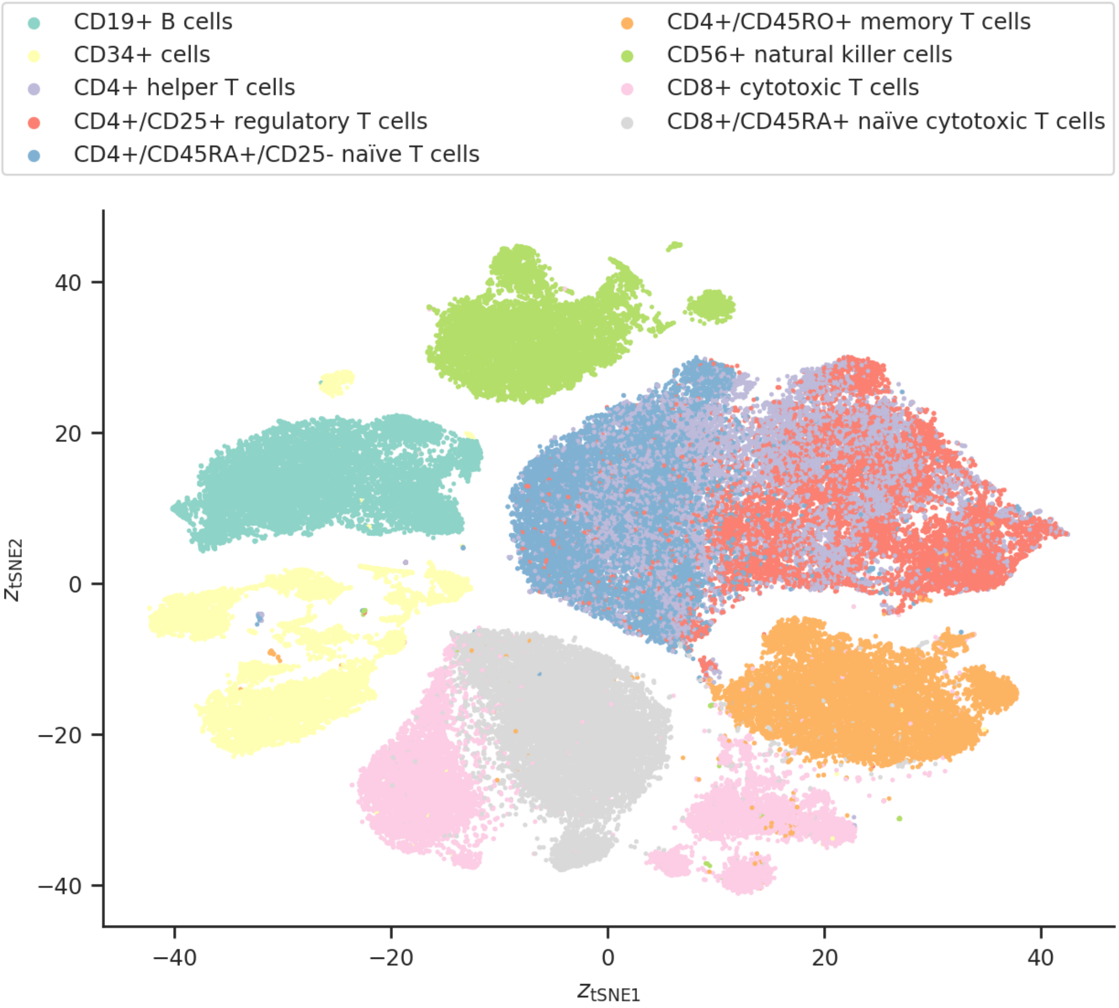


**Figure S10.** Latent space of the median-performing GMVAE-NB model both trained and evaluated on the full PBMC data set. The encoding of the cells in the latent space has been embedded in two dimensions using *t*-SNE and are colour-coded using their cell types. Clear separation can be seen corresponding to different cell types, and some similar cell types are also clustered close together or mixing together.

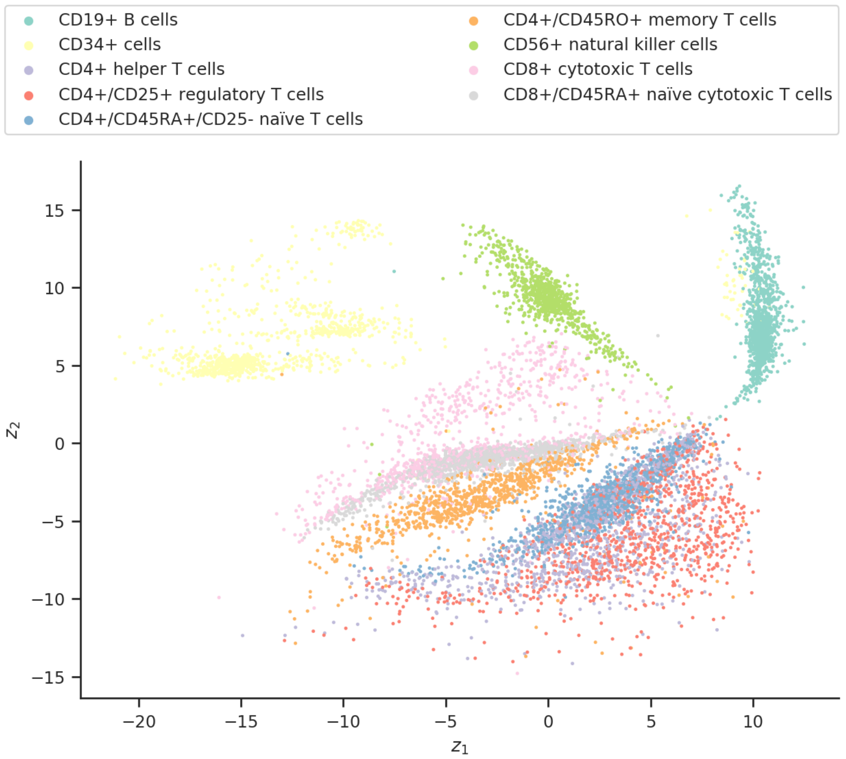


**(a)**

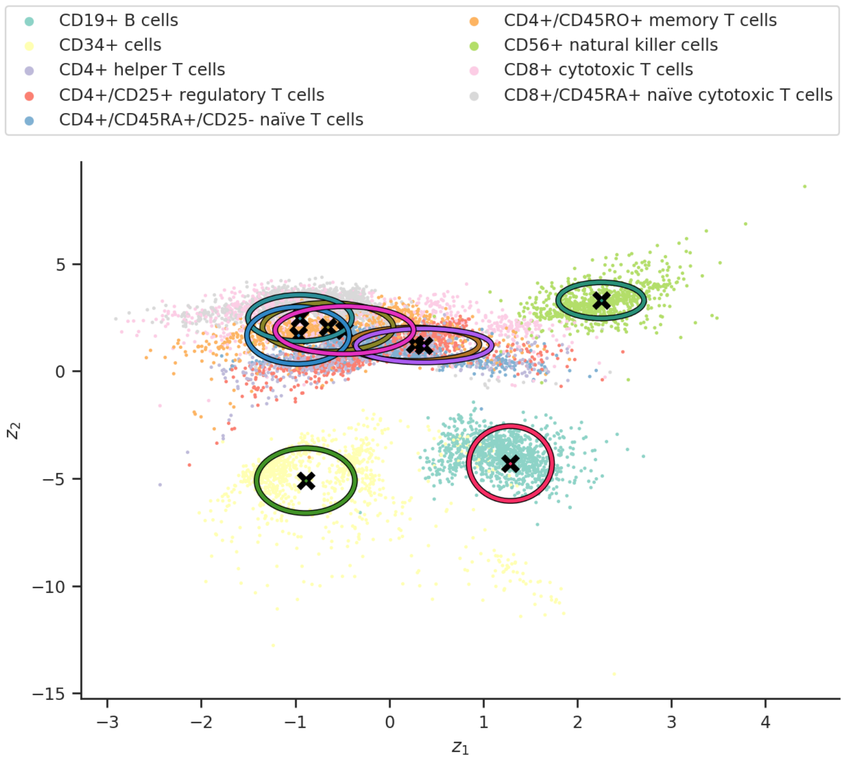


**(b)**

**Figure S11.** Latent space of **(a)** the scvis model and **(b)** the median-performing GMVAE-NB model with a 2-d latent variable (the mean and the covariance of each component of $p_{\boldsymbol{\theta}}(\mathbf{z} \mid y)$ of the GMVAE is also plotted) trained and evaluated on the PBMC data set. The encoding of the cells in the latent space has been embedded in two dimensions using $t$-SNE and are colour-coded using their cell types.

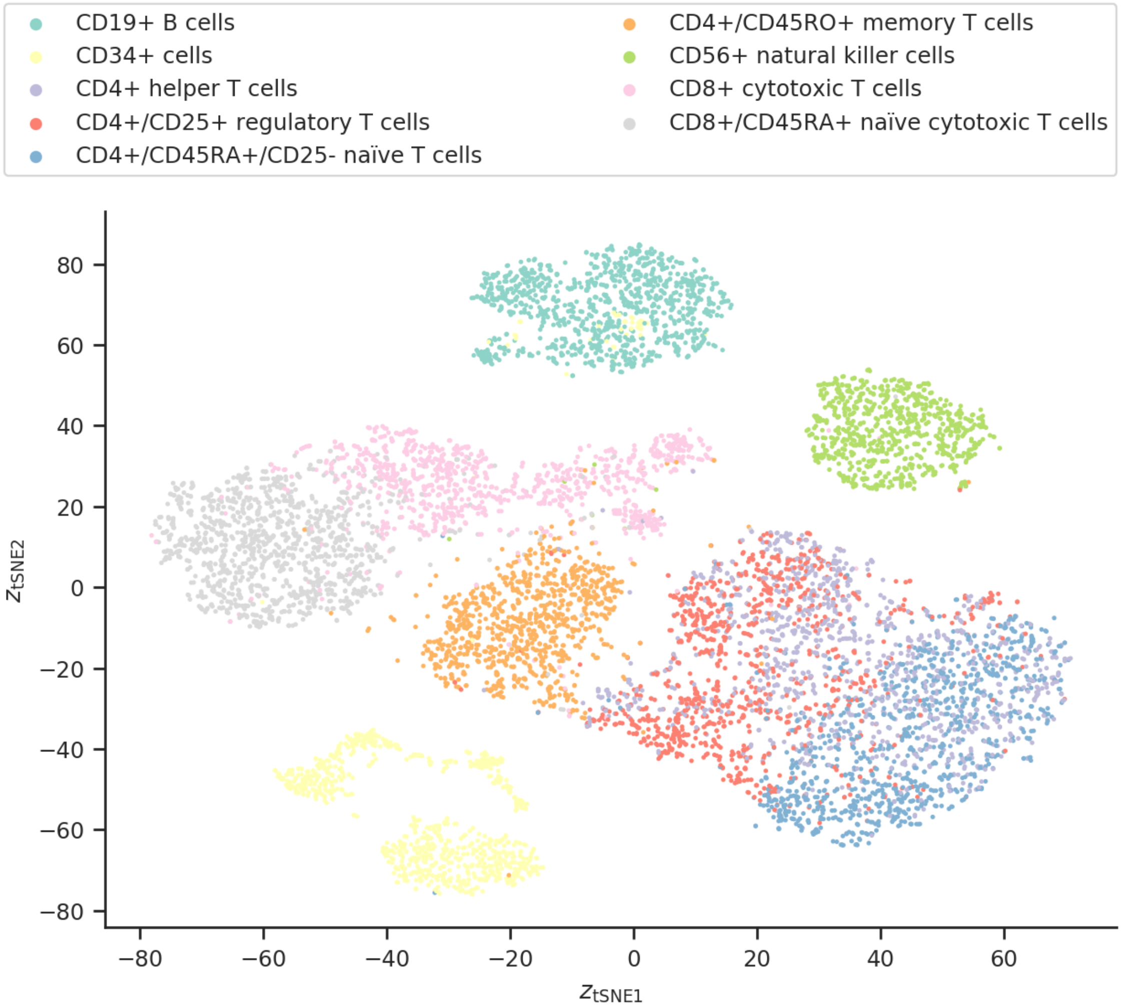


**Figure S12.** Latent space of the median-performing scVI model trained and evaluated on the PBMC data set. The encoding of the cells in the latent space has been embedded in two dimensions using *t*-SNE and are colour-coded using their cell types.

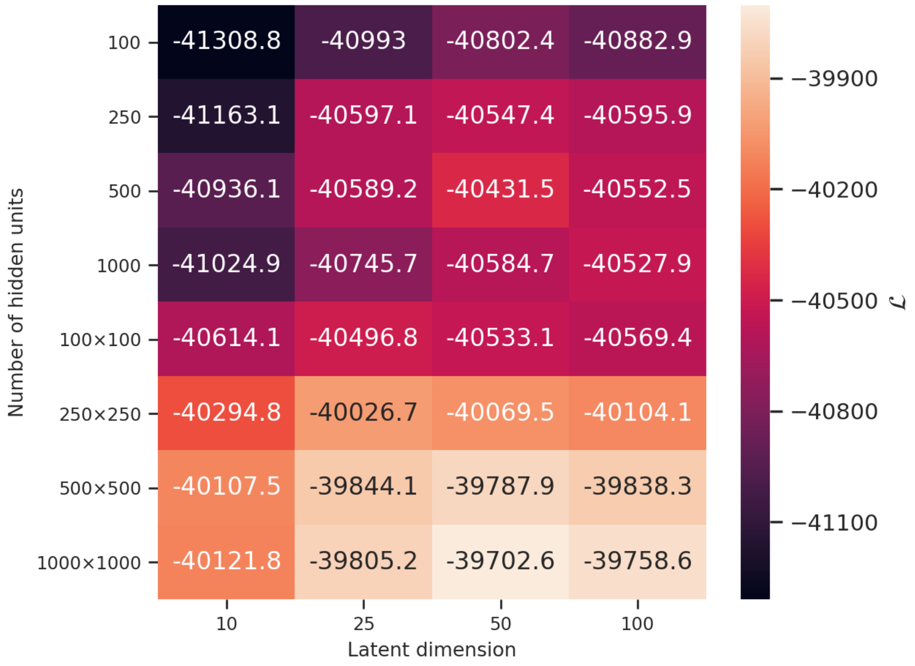


**Figure S13.** Test marginal log-likelihood lower bound, $\mathcal{L}$, for the gene-name-mapped TCGA data set limited to the 5000 most varying genes using a VAE-NB model with different latent dimensions and different number of hidden units. The multiplication sign ($\times$) separate the number of units in each hidden layer. The most optimal architecture is two hidden layers of 1000 units each and 50 latent dimensions.

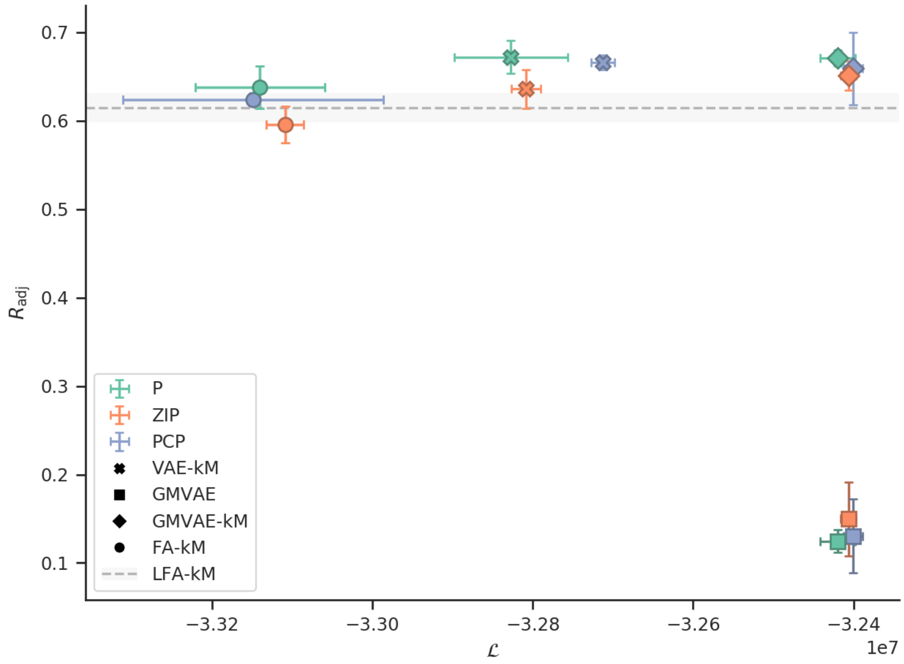


**(a)**

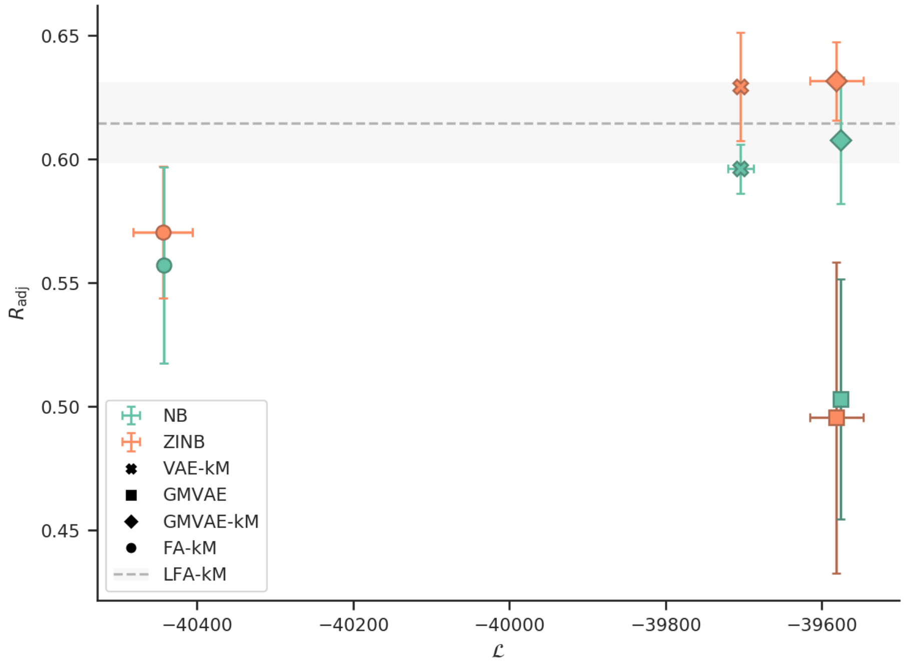


**(b)**

**Figure S14.** Comparison of VAE, GMVAE, and FA models with **(a)** Poisson and **(b)** negative-binomial likelihood functions trained and evaluated on the gene-name-mapped TCGA data set limited to the 5000 most varying genes. For each combination the mean and the standard deviation of the adjusted Rand index are plotted against the marginal log-likelihood lower bound. Note the difference in lower bound between the two figures. The mean and the standard deviation of the adjusted Rand index are also plotted for the LFA baseline model.

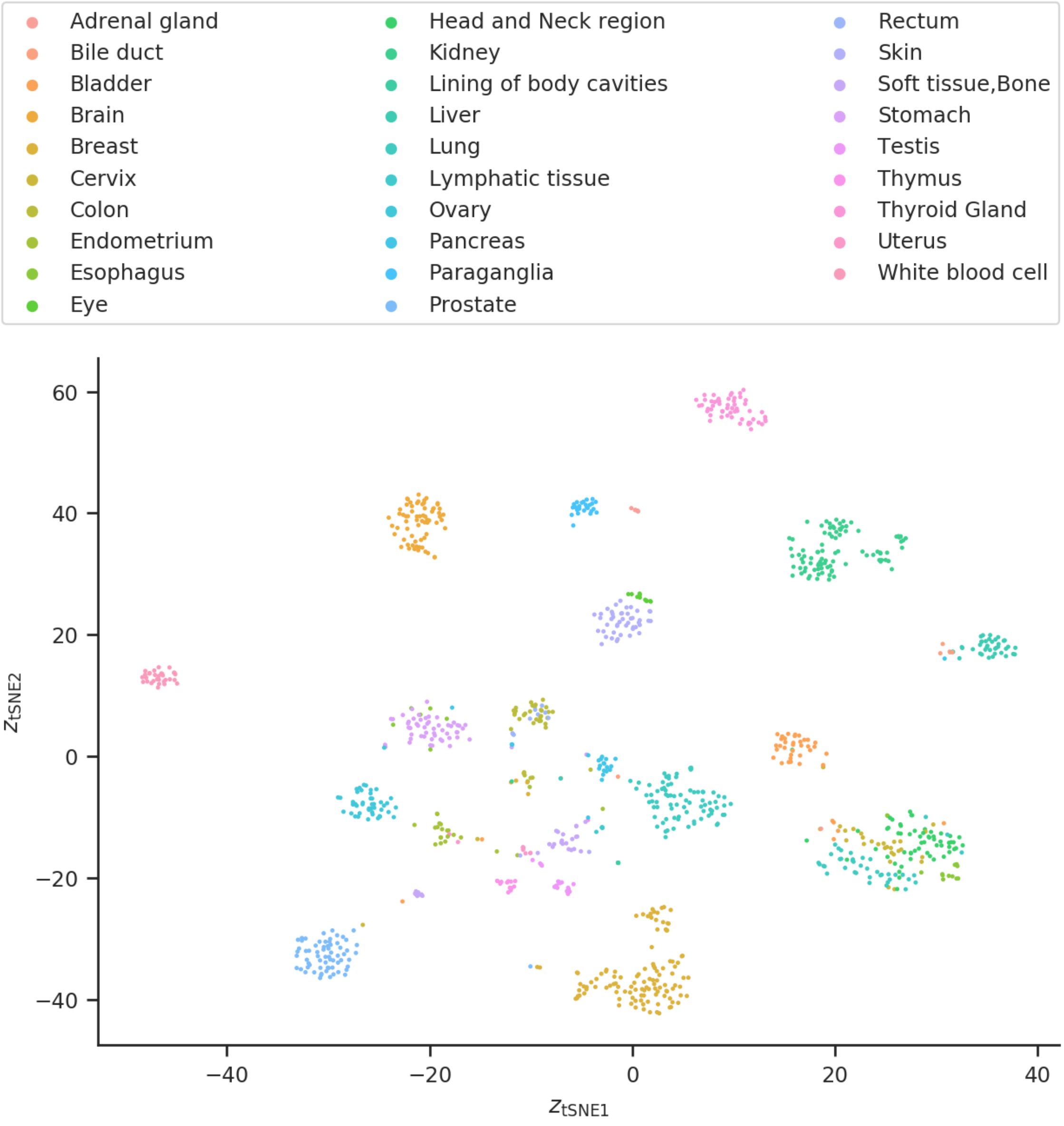


**Figure S15.** Latent space of the median-performing GMVAE-NB model trained and evaluated on the gene-name-mapped TCGA data set limited to the 5000 most varying genes. The encoding of the cells in the latent space has been embedded in two dimensions using *t*-SNE and are colour-coded using the tissue sites to which they belong. Many of the clusters are clearly separated and they correspond to different tissue sites.

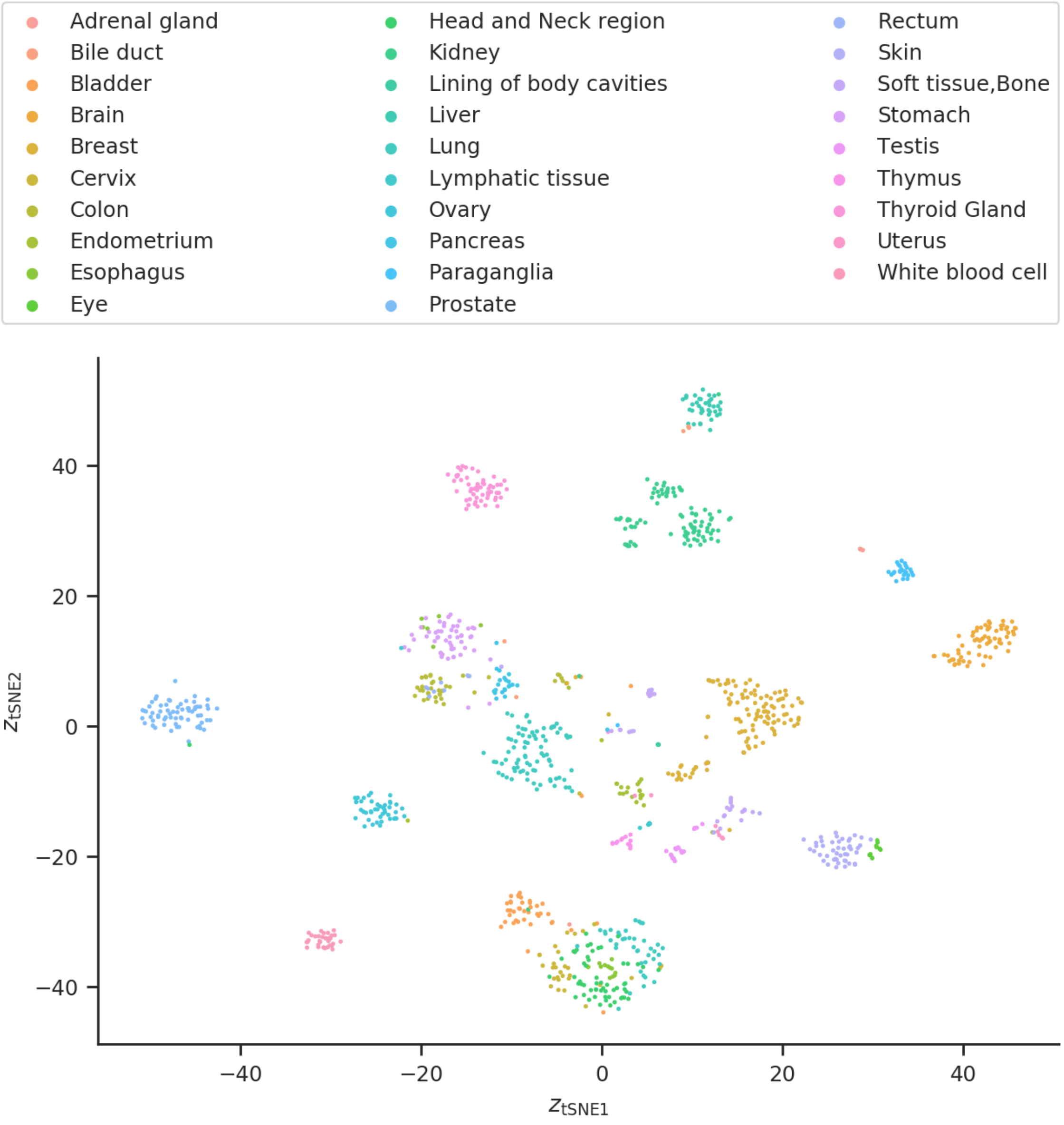


**Figure S16.** Latent space of the median-performing VAE-NB model trained and evaluated on the gene-name-mapped TCGA data set limited to the 5000 most varying genes. The encoding of the cells in the latent space has been embedded in two dimensions using *t*-SNE and are colour-coded using the tissue sites to which they belong. Many of the clusters are clearly separated and they correspond to different tissue sites.

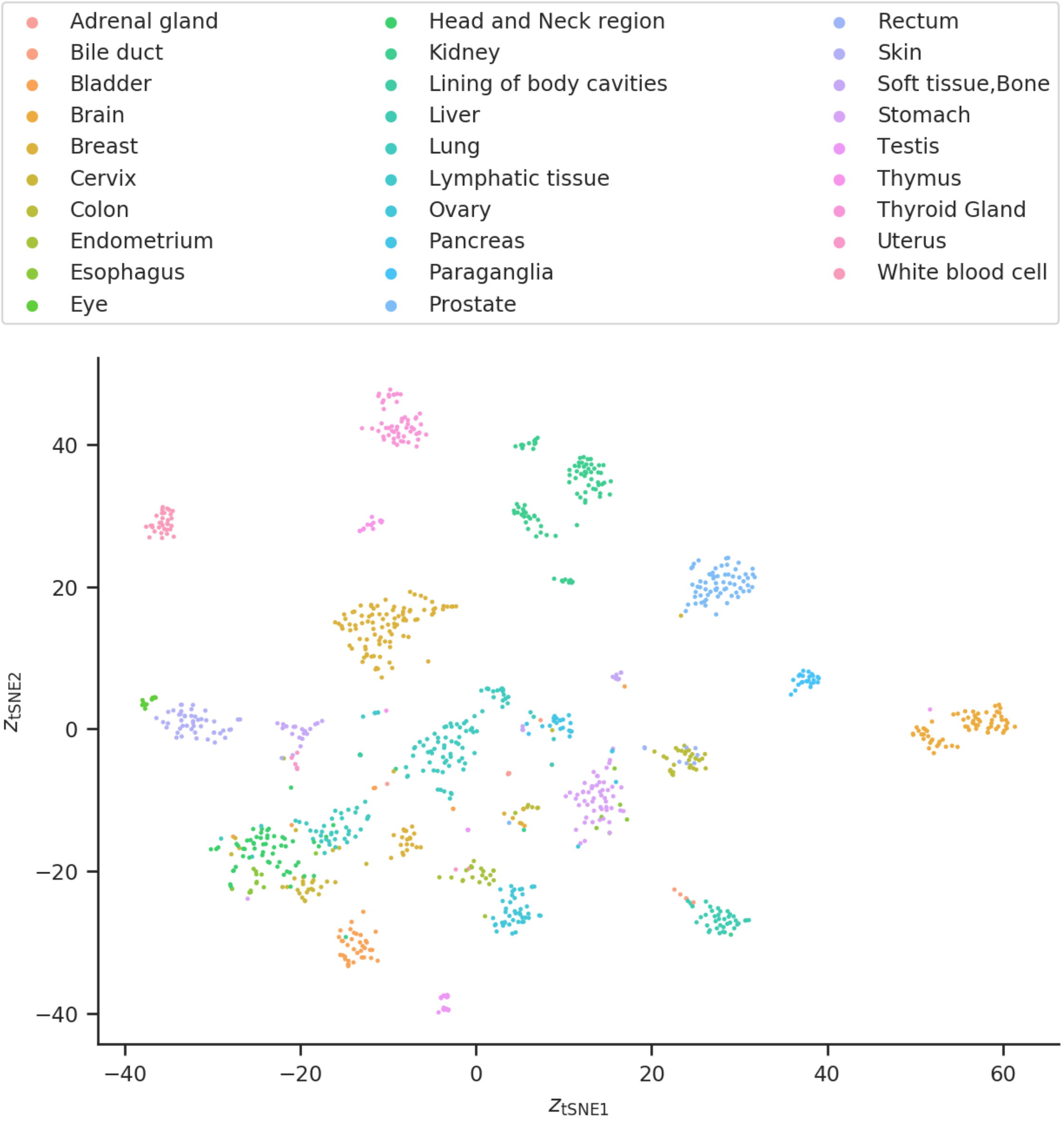


**Figure S17.** Latent space of the median-performing FA-NB model trained and evaluated on the gene-name-mapped TCGA data set limited to the 5000 most varying genes. The encoding of the cells in the latent space has been embedded in two dimensions using *t*-SNE and are colour-coded using the tissue sites to which they belong. Many of the clusters are clearly separated and they correspond to different tissue sites.

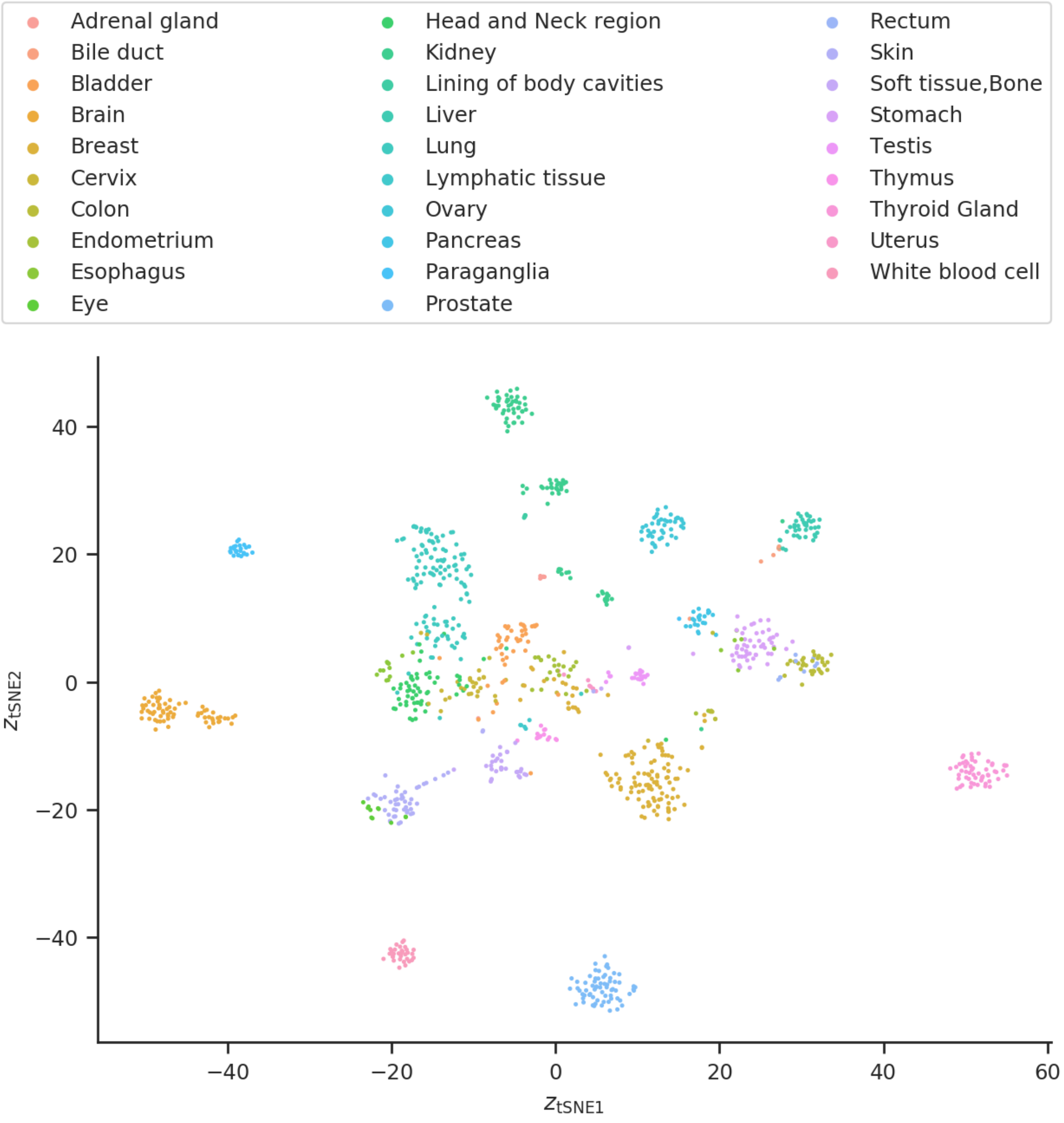


**Figure S18.** Latent space of the median-performing LFA baseline model trained and evaluated on the gene-name-mapped TCGA data set limited to the 5000 most varying genes. The encoding of the cells in the latent space has been embedded in two dimensions using *t*-SNE and are colour-coded using the tissue sites to which they belong. Many of the clusters are clearly separated and they correspond to different tissue sites.

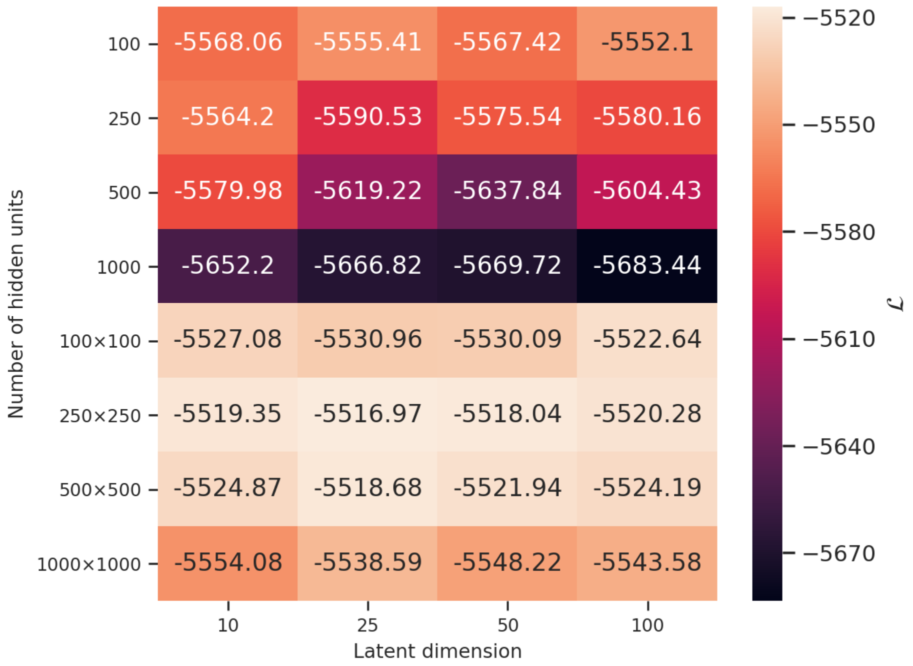


**Figure S19.** Test marginal log-likelihood lower bounds, $\mathcal{L}$, for the MBC-20k data set using a VAE-NB model with different latent dimensions and different number of hidden units. The multiplication sign $(\times)$ separate the number of units in each hidden layer. The most optimal architecture is two hidden layers of 250 units each and 25 latent dimensions.

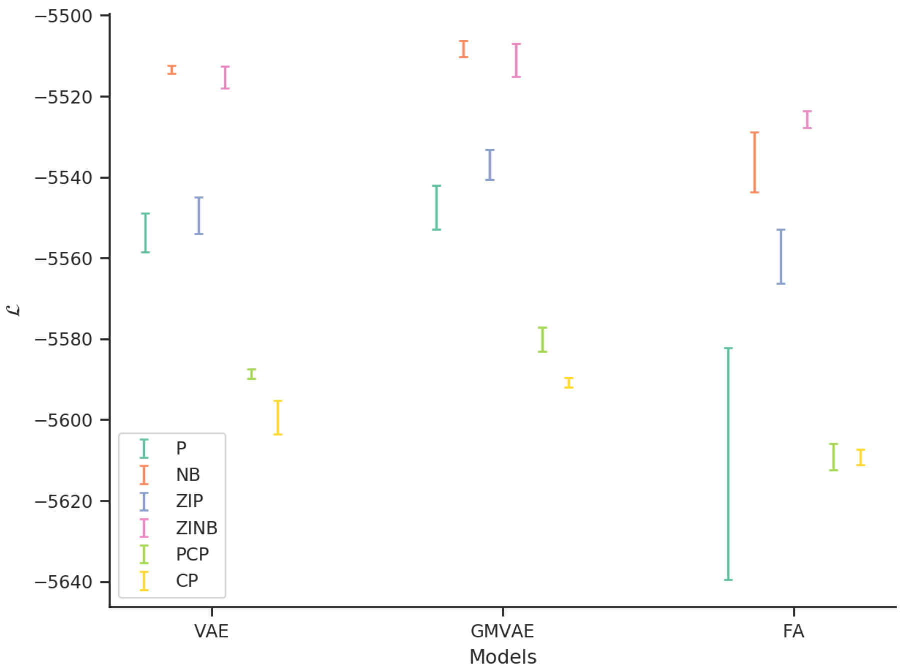


**Figure S20.** Comparison of VAE, GMVAE, and FA models with different likelihood functions trained and evaluated on the MBC-20k data set. For each combination the mean and the standard deviation of the marginal log-likelihood lower bound is plotted.

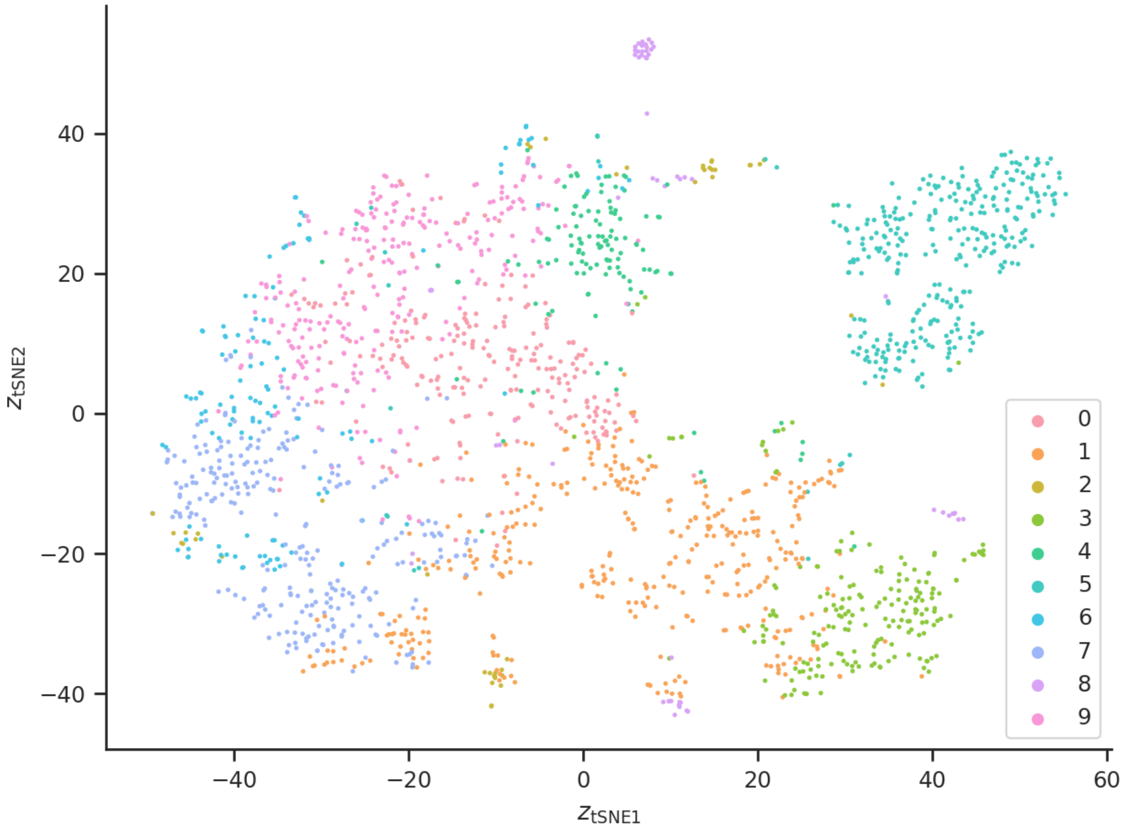


**Figure S21.** Latent space of the GMVAE-NB trained and evaluated on the MBC-20k data set. Ten clusters was used in the model, and the encoding of the cells in the latent space has been embedded in two dimensions using *t*-SNE and are colour-coded using the clusters to which they belong. Different regions can be seen with even some separation for some clusters.

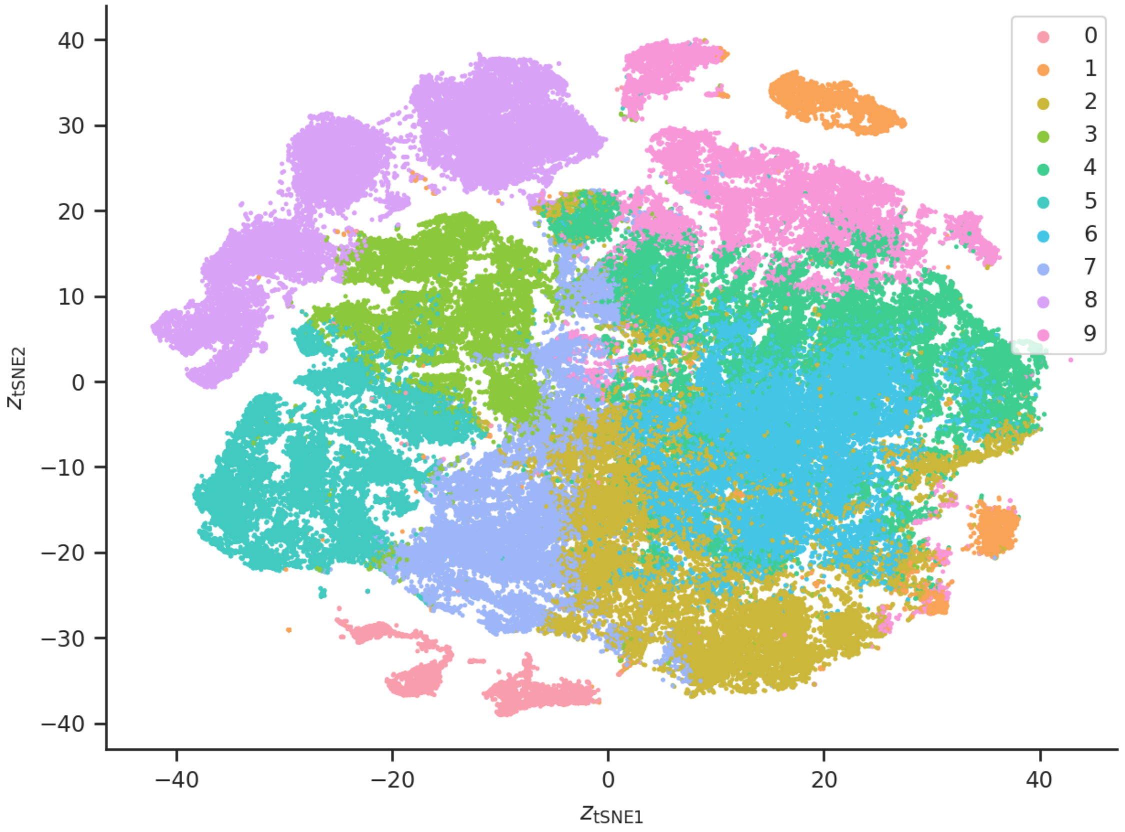


**Figure S22.** Latent space of the GMVAE-ZINB model trained and evaluated on the MBC data set. Ten clusters was used in the model, and the encoding of the cells in the latent space has been embedded in two dimensions using *t*-SNE and are colour-coded using the clusters to which they belong. Different regions can be seen with even some separation for some clusters.

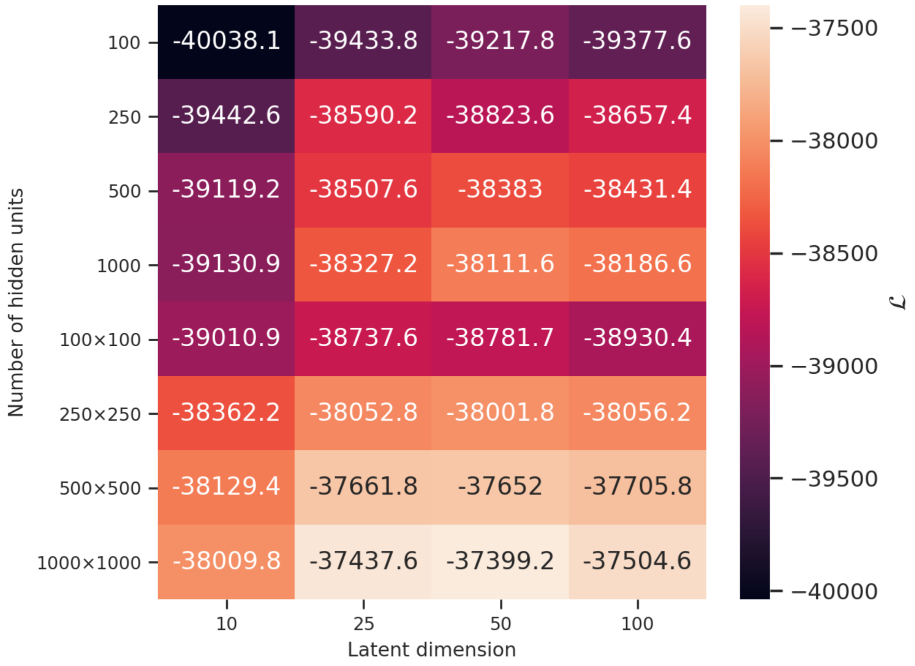


**Figure S23.** Test marginal log-likelihood lower bounds, $\mathcal{L}$, for the gene-name-mapped GTEx data set limited to the 5000 most varying genes using a VAE-NB model with different latent dimensions and different number of hidden units. The multiplication sign ($\times$) separate the number of units in each hidden layer. The most optimal architecture is two hidden layers of 1000 units each and 50 latent dimensions.

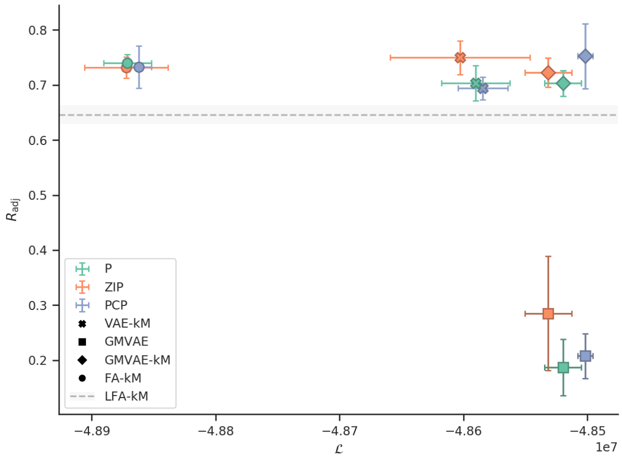


**(a)**

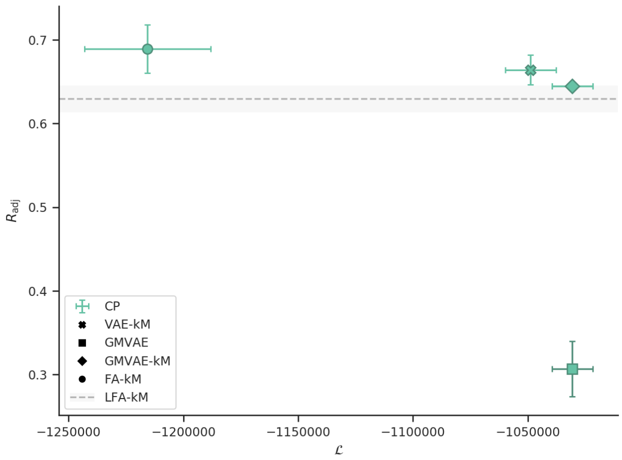


**(b)**

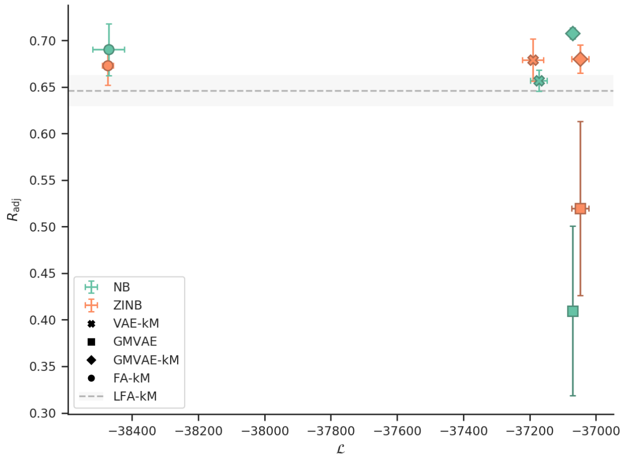


**(c)**

**Figure S24.** Comparison of VAE, GMVAE, and FA models with **(a)** Poisson, **(b)** constrained Poisson, and **(c)** negative-binomial likelihood functions trained and evaluated on the gene-name-mapped GTEx data set limited to the 5000 most varying genes. For each combination the mean and the standard deviation of the adjusted Rand index are plotted against the marginal log-likelihood lower bound. Note the difference in lower bound between the figures. The mean and the standard deviation of the adjusted Rand index are also plotted for the LFA baseline model.

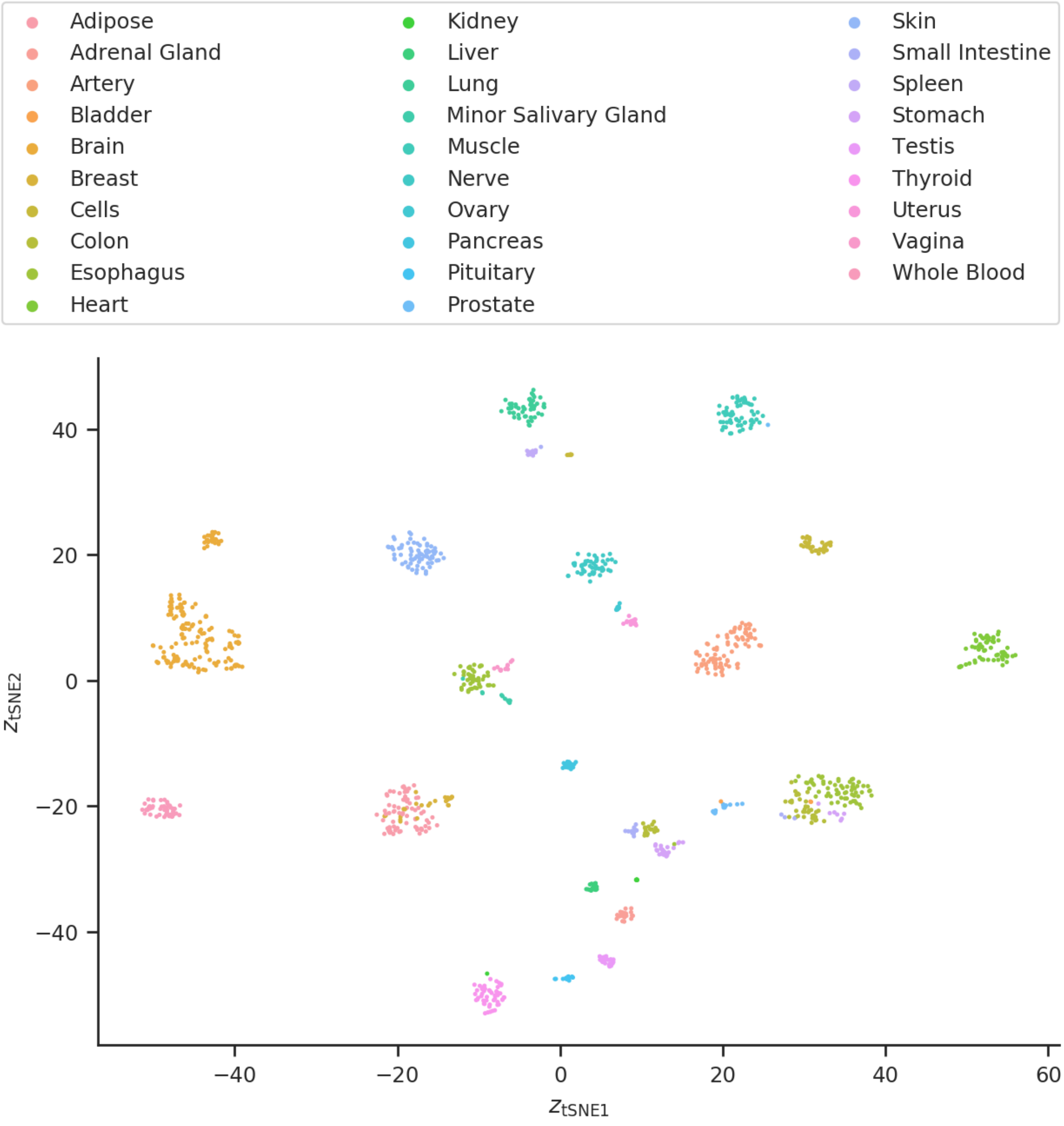


**Figure S25.** Latent space of the median-performing GMVAE-ZINB trained and evaluated on the gene-name-mapped GTEx data set limited to the 5000 most varying genes. The encoding of the cells in the latent space has been embedded in two dimensions using *t*-SNE and are colour-coded using the tissue sites to which they belong. Many of the clusters are clearly separated and they correspond to different tissue sites.
